# Identification and molecular characterization of a novel TYLCV isolate breaking bred-resistance to threaten tomato cultivar

**DOI:** 10.64898/2026.06.16.732612

**Authors:** Yaogui Zhou, Shihuai Jin, Jing Zhong, Xingming Xiao, Ming Ding, Liling Zhao, Zhongxin Guo

## Abstract

Tomato yellow leaf curl virus (TYLCV) is a devastating viral pathogen threatening agricultural crops globally. In this study, we identified a novel TYLCV isolate (TYLCV-YN6244), which caused viral epidemic in resistant tomato cultivars at Yuanmo county, Yunnan Province of China. We determined the complete genome of TYLCV-YN6244 and found it encoded six viral proteins characteristic of *Geminivirus*. We identified its V2 protein as a potent viral suppressor of RNA silencing (VSR), and generated infectious clone of wildtype TYLCV-YN6244, or V2-defective TYLCV-YN6244 (TYLCV-YN6244-ΔV2) in which *V2* was deleted. Both of infectious clones were capable of systemically infecting tobacco and tomato. However, TYLCV-YN6244 but not TYLCV-YN6244-ΔV2 could cause disease symptoms in wildtype tobacco or tomato plants, and viral accumulation was drastically reduced in plants infected with TYLCV-YN6244-ΔV2 compared to TYLCV-YN6244 while the efficiency of virus-derived small interfering RNAs (vsiRNAs) biogenesis was conversely increased in plants infected with TYLCV-YN6244-ΔV2. Surprisingly, small RNA profiling indicated that 21nt and 22nt rather than 24nt vsiRNAs were predominantly produced in tomato plants infected with either TYLCV-YN6244 or TYLCV-YN6244-ΔV2. Furthermore, transcriptome analyses revealed that TYLCV-YN6244 or TYLCV-YN6244-ΔV2 infection differentially modulated metabolism and defense-related pathways in tomato, probably underlying distinct viral pathogenicity and disease symptoms induced in plants. Overall, our research not only identified a novel pathogenic TYLCV isolate but also characterized molecular biology and host response in tomato with infectious clones firstly developed, with implications in untangling virus-host interaction for developing novel resistance in crop tomato.

## Background

Tomato yellow leaf curl virus (TYLCV), a member of the genus *Begomovirus* in the family *Geminiviridae*, is a species of plant DNA virus causing disastrous diseases in many economically important crops, including tomato ^[1]^. TYLCV contains a single circular ssDNA genome, DNA-A, which encodes six typical Open Reading Frames (ORFs): V1, V2, C1, C2, C3, and C4, being organized in two transcriptional directions separated by an intergenic region (IR) ^[2]^. While C1 and C3 are associated with viral replication, C2 acts as a transcriptional activator involving in gene activation and C4 is involved in disease induction with VSR activities ^[3, 4]^. V2 also possesses VSR activities to suppress RNA silencing ^[5,6]^, whereas V1 is the coat protein (CP) packaging the viral genome into an icosahedral particle and functions in insect transmission and long-distance virus movement ^[7]^. Recently, several small proteins encoded by geminiviruses have also been identified and characterized to be implicated in host-virus interactions ^[8–10]^. TYLCV is mainly transmitted by insect vectors whitefly *Bemisia tabaci* in a persistent-circulative or persistent-propagative manner ^[11,12]^ and spread to cause epidemics of Tomato yellow leaf curl disease (TYLCD) ^[2]^. Since the first discovery of TYLCD in the late 1930s, TYLCD remains to be one of the top ten viral diseases threatening tomato production globally ^[2]^.

Breeding resistant germplasm is the most cost-effective and ecologically friendly approach to control viral diseases in agricultural production ^[13,14]^. Considerable efforts have been made to identify resistance genes for breeding TYLCV-resistance tomato cultivars. So far, six different resistance genes (*Ty-1*, *Ty-2*, *Ty-3*, *Ty-4*, *ty-5*, and *Ty-6*) have been identified from wild tomato species, including *S. habrochaites* and *S. chilense* ^[15–19]^. *Ty-1* and *Ty-3*, which are allelic, encode an RNA-dependent RNA polymerase and belong to the RDRγ-like protein family ^[16]^. *Ty-2* encodes a nucleotide-binding leucine-rich repeat protein (NLR) that recognizes the C1 protein to activate resistance ^[18,20]^. *Ty-1*, *Ty-2*, and *Ty-3* have been widely applied for breeding resistant tomato cultivars in the past decade ^[18]^. However, new TYLCV isolates emerge to break bred-resistance from time to time to cause epidemics in agricultural production ^[21, 22]^.

RNA interference (RNAi)-based antiviral defense is an evolutionarily conserved mechanism counteracting the invasion of all different RNA or DNA viruses in eukaryotes, showing significant potential for developing broad-spectrum antiviral resistance ^[23, 24]^. In model plant Arabidopsis, Dicer-like enzymes (DCLs) perceive and dice viral double-stranded RNA (dsRNA) of replication intermediates to produce primary vsiRNAs and initiate the antiviral defense. Upon the infection of RNA viruses, DCL4 and DCL2 perceive viral dsRNAs and respectively produces the 21 or 22 nucleotides (nt) primary vsiRNAs to initiate antiviral defense ^[25]^. RNA-dependent RNA polymerases 1 (RDR1) and RDR6, together with antiviral RNAi-defective 1 (AVI1) and AVI2, further amplify viral dsRNAs to produce secondary vsiRNAs ^[26, 27]^. Primary and secondary 21 or 22nt vsiRNAs are loaded into Argonaute (AGO) proteins (preferably AGO1 and AGO2) to form the RNA-induced silencing complex (RISC), then mediates antiviral RNAi through post-transcriptional gene silencing (PTGS) pathway ^[28]^. Upon the infection of DNA viruses, DCL3 and RDR2 produce 24nt primary and secondary vsiRNAs, which are incorporated into AGO4, AGO6, or AGO9 to induce resistance mainly through RNA-directed DNA methylation (RdDM) mediated transcriptional gene silencing (TGS) pathway ^[29]^.

However, during long-term arms race between viruses and plants, viruses evolve different Viral Suppressor of RNAi (VSR) to repress antiviral RNAi and function as determinant factor of viral virulence and disease induction in host plants ^[30, 31]^. Nearly all known pathogenic plant viruses encode at least one VSR, the prevalence of VSRs hinders our appreciation of antiviral RNAi immunity and has been neck-bottled research in the field. Interestingly, a sensitized forward genetic screen was recently developed to dissect the antiviral immunity in Arabidopsis and tomato plants, based on the VSR 2b-deficient cucumber mosaic virus (CMV-Δ2b) ^[32, 33]^. However, the small RNA-based antiviral immunity is still not clear, requesting further elucidation to be translated for developing applicable resistance in plants including agricultural crops ^[34, 35]^. Notably, a comparable genetic system has not been well established to uncover novel components for elucidating the antiviral defense to plant DNA viruses.

In this study, we identified a novel TYLCV isolate, TYLCV-YN6244 (Accession MN503102), that causing severe disease in field-grown tomato cultivars with *Ty-1*/*Ty-2*-mediated resistance at Yuanmou county, Yunnan Province of China. After determining the activity of its VSR V2 protein, we developed infectious clones of wildtype TYLCV-YN6244 or V2-deficient TYLCV-YN6244 (TYLCV-YN6244-ΔV2). We demonstrated that both infectious clones were also able to systemically infect plants. However, in contrast to TYLCV-YN6244-ΔV2, TYLCV-YN6244 caused severe disease symptoms in plants, and the virus was accumulated remarkably more while the efficiency of vsiRNA biogenesis was dramatically decreased, indicating the crucial roles of V2 in suppression of antiviral RNAi and disease induction in host plants. Surprisingly, small RNA-seq analyses revealed that 21nt and 22nt rather than 24nt vsiRNAs were predominantly produced with the infection of TYLCV-YN6244 or TYLCV-YN6244-ΔV2, suggesting that 21nt and 22nt vsiRNAs may also mediate antiviral RNAi in defense of the infection of plant DNA viruses. Furthermore, transcriptome profiling through RNA-seq showed that TYLCV-YN6244 or TYLCV-YN6244-ΔV2 infection differentially modulated metabolism and defense pathways in tomato, which probably contributes to the differences in the viral accumulation and the disease symptoms induced in plants. Thus, this study not only identified a novel pathogenic TYLCV isolate but also characterized molecular biology and host responses, providing new resources and knowledge for studying host-virus interaction, especially antiviral RNAi immunity to TYLCV, an important plant DNA virus, for breeding new resistance in crop plants.

## Methods

### Plant materials and growth conditions

Tomato plants showing disease symptoms of TYLCD were collected from Yuanmou, Yunnan Province of China. Tobacco (*N. benthamiana*), tomato (Micro-Tom) plants used for virus inoculation were grown in an insect-free growth room at 24 °C under 8 h light/16 h dark cycle.

### Phylogenetic analysis

The complete genome sequence of TYLCV-YN6244 was aligned with known viruses deposited in the GenBank database and a phylogenetic tree was constructed through MEGA11 by neighbor-joining method with 1000 bootstrap replicates.

### Construction of transient expression vectors

According to the method of homologous recombination, we designed primers with restriction sites and homology arms to amplify full-length genes of *C1*, *C2*, *C3*, *C4*, *C5*, *V1*, *V2*, *V3*. The amplified PCR products were cloned into the pCAMBIA3301 vector and confirmed by Sanger sequencing to produce pCAMBIA3301-flag-C1, pCAMBIA3301-flag-C2, pCAMBIA3301-flag-C3, pCAMBIA3301-flag-C4, pCAMBIA3301-flag-C5, pCAMBIA3301-flag-V1, pCAMBIA3301-flag-V2, and pCAMBIA3301-flag-V3. Subsequently, each recombinant vector was transformed by heat shock method into *Agrobacterium tumefaciens* strain *GV3101*. Primers used in this study are listed in Supplementary Table S5.

### Assay of VSR activity

Agrobacterium transformed with pCAMBIA3301-flag-C1, pCAMBIA3301-flag-C2,pCAMBIA3301-flag-C4,pCAMBIA3301-flag-C5,pCAMBIA3301-flag-V1,pCAMBIA3301-flag-V2,pCAMBIA3301-flag-V3, pCAMBIA3301, pGD-GFP and pGD-VSRs were cultured in lysogeny broth fluid medium at 28 °C, pelleted and resuspended in a buffer containing 10 mM MES, 10 mM MgCl_2_, 150 mM acetosyringone. Agrobacterium solutions were diluted to OD_600_ = 0.8. pGD-GFP was combined with viral genes or pGD-VSRs and co-infiltrated into leaves of 16c transgenic *N. benthamiana* plants using 1 mL needleless syringes as described ^[36, 37]^. Vector pGD-VSRs (embedded with P19, HcPro and γb) was used for positive control of VSR ^[38]^, empty vector for negative control. GFP signal was imaged with 488-nm excitation and a detection window of 495 to 535 nm under Zeiss LSM880 confocal microscope.

### Construction of the infectious clone of TYLCV-YN6244 and TYLCV-YN6244-**Δ**V2

To construct the infectious clone of TYLCV-YN6244 and TYLCV-YN6244-ΔV2, the full-length and *V2*-mutated (1-30 aa deleted) fragment, in which V2 domain but no other overlapped genes was mutated, of 0.5-unit DNA A was amplified using primers TYLCV-YN6244-0.5-unit-F/R or TYLCV-YN6244-ΔV2-0.5-unit-F/R, and cloned into pGEM-Teasy (A1360, Promega, Madison, USA) to produce pGEM-Teasy-TYLCV-YN6244-0.5-unit DNA and pGEM-Teasy-TYLCV-YN6244-ΔV2-0.5-unit. The correct clone pGEM-Teasy-TYLCV-YN6244-0.5-unit or pGEM-Teasy-TYLCV-YN6244-ΔV2-0.5-unit DNA were digested with *Sac*lJ (R3156V, NEB, Lpswich, USA) and *Hind*lJ (R0104V, NEB, Lpswich, USA) and introduced into the binary vector pBinplus (V010906, NovoPro, Shanhai, China) to produce pBinplus-TYLCV-YN6244-0.5-unit or pBinplus-TYLCV-YN6244-ΔV2-0.5-unit DNA. Subsequently the full-length or V2 mutant fragment of 1-unit DNA was amplified using primers TYLCV-YN6244-1-unit-F/R or TYLCV-YN6244-ΔV2-1-unit-F/R, and cloned into pEASYTM-T5-Zero (CT501-01, TransGen, Beijing, China) to produce pEASYTM-T5-Zero-TYLCV-YN6244-1-unit or pEASYTM-T5-Zero-TYLCV-YN6244-ΔV2-1-unit DNA. The correct clone pEASYTM-T5-Zero-TYLCV-YN6244-1-unit or pEASYTM-T5-Zero-TYLCV-YN6244-ΔV2-1-unit DNA were digested with *Sac*lJ and introduced into pBinplus-TYLCV-YN6244-0.5-unit or pBinplus-TYLCV-YN6244-ΔV2-0.5-unit DNA to produce pBinplus-TYLCV-YN6244-1.5-unit or pBinplus-TYLCV-YN6244-ΔV2-1.5-unit vector.

All vectors were sequenced. The correct pBinplus-TYLCV-YN6244-1.5-unit or pBinplus-TYLCV-YN6244-ΔV2-1.5-unit vector were transformed into the *Agrobacterium tumefaciens* strain *GV3101* by heat shock method. Primers used in this study are listed in Supplementary Table S5.

### Agrobacterium-mediated inoculation of infectious clone to tobacco and tomato plants

Agrobacterium harboring TYLCV-YN6244 and TYLCV-YN6244-ΔV2 were cultured in lysogeny broth fluid medium at 28 °C, pelleted and resuspended in a buffer containing 10 mM MES, 10 mM MgCl_2_, 150mM acetosyringone. The Agrobacterium solutions were diluted to OD_600_ = 0.8, and infiltrated into leaves of tobacco and tomato plants at the growth stage with only two cotyledons, using 1 mL needleless syringes as described ^[39]^.

### Protein extraction and Western blot analysis

At 15-day post inoculated by TYLCV-YN6244 or TYLCV-YN6244-ΔV2, total proteins were extracted from leaf samples according to a method described in published research ^[40]^. Leaves from 3 individual plants were pooled for each biological replicate. Equal amounts of proteins were transferred to PVDF membranes after being separated on 10 or 12.5% SDS–PAGE gels. GFP-fused proteins were detected using rabbit monoclonal anti-GFP antibody (1:3000; ab316291, Abcam, Cambridge, UK). The blot signal was detected with a chemiluminescence image analysis system (Tanon-5200).

### DNA extraction and Southern blot analysis

At 21-day post inoculated by TYLCV-YN6244 or TYLCV-YN6244-ΔV2, the upper systemic leaves were collected for extracting DNA. Leaves from 3 individual plants were pooled for each biological replicate. DNA extraction and Southern blots were conducted according to a published protocol ^[37]^. Southern blots were performed with probes of Biotin-11-dUTP (R0081, Thermos Scientific, Waltham, USA) labeled cDNA or DNA oligonucleotides, as described in previous research ^[41]^. The blot signal was detected with a chemiluminescence image analysis system (Tanon-5200). Probe primers used in this study are listed in Supplementary Table S5.

### RNA extraction and Northern blot analysis

At 21-day post inoculated by TYLCV-YN6244 or TYLCV-YN6244-ΔV2, RNA was extracted from the upper systemic leaves of plants. Leaves from 3 individual plants were pooled for each biological replicate. Total and small RNAs were isolated using the hot phenol method, and Northern blot was performed as described previously ^[27]^, with biotin-labeled DNA probes using Biotin-11-dUTP. The blot signal was detected with a chemiluminescence image analysis system (Tanon-5200). Probe primers used in this study are listed in Supplementary Table S5.

### Real-time quantitative PCR

Accumulations of genomic DNA of TYLCV-YN6244 or TYLCV-YN6244-ΔV2 were examined by absolute real-time quantitative PCR. Plasmid of TYLCV-YN6244 was used be standard sample. Reactions were performed using the Taq Pro Universal SYBR qPCR Master Mix (Q172, Vazyme, Nanjin, China) with specific primers listed in Supplementary Table S5.

### RNA-Seq and small RNAs-Seq

Paired-end mRNA libraries were generated using NEBNext® UltraTM RNA Library Prep Kit for Illumina® (E7770, NEB, Lpswich, USA) according to the manufacturer’s manual and were sequenced on an Illumina HiSeq 4000 platform; 150bp paired-end reads were generated. sRNA libraries were generated using NEBNext® Multiplex Small RNA Library Prep Set for Illumina® (E7560S, NEB, Lpswich, USA.) and sequenced on an Illumina HiSeq2500 platform, and 51bp single-end reads were generated.

### Differential expression analysis

Raw counts were used for differential expression analysis using DESeq2 v.1.32.0 to obtain DESeq2-normalized counts through the median of ratios method ^[42]^. The DESeq2 design matrix contained information for tomato infected with Mock, TYLCV-YN6244 or TYLCV-YN6244- Δ V2 (simplified as MT-Mock, MT-TYLCV-YN6244, MT-TYLCV-YN6244-ΔV2).

### Gene ontology (GO) enrichment analysis and venn analysis

Gene Ontology (GO) annotations of the contigs were determined using ClusterProfile ^[42]^. GO-enrichment analysis of DEGs was performed using the ClusterProfile R pock ^[43]^, and the Padjust-value were calculated using the FalseDiscovery Rate (FDR) correction. Compared to Mock plants, genes with Padjust-value < 0.05 were included in our analysis. The overlapping induced or suppressed genes were analyzed.

### Small RNA analysis

The quality of clean reads was also checked by FastQC. The adapter sequence was removed using cutadapt, and clean reads of small RNAs with size from 18 to 26nt were extracted. Subsequently, FASTA data were aligned to TYLCV-YN6244 genome using Bowtie (v1.3.1) to identify vsiRNA reads. vsiRNA and remaining small RNA reads with perfect matching were then respectively mapped to the virus genome or host genome for profiling analysis.

## Results

### Identification of a novel pathogenic TYLCV isolate in tomato cultivars

During a survey of cultivated tomato-infecting TYLCV, in agricultural fields at Yuanmou county, Yunnan Province of China, we observed that tomato varieties, including Zuanhong No.5 and Baxi which are bred with *Ty-1* and *Ty-2*-mediated resistance to TYLCV, displayed typical symptoms of TYLCD, such as leaf curling, chlorosis, and stunting (Fig. 1A; Supplementary Fig. S1). The complete nucleotide sequence of the TYLCV-Yunnan isolate (TYLCV-YN6244) was determined to be 2781nt in length (Supplementary Table S1) and contains six typical ORFs (V1, V2, C1, C2, C3 and C4) and two small ORFs (V3 and C5) (Fig. 1B; Supplementary Table S2). These ORFs respectively encode viral replication, coat protein (CP), movement protein (MP), VSRs, and other small proteins that play roles in various stages of the virus life cycle. Interestingly, alignment of TYLCV-YN6244 genome to known TYLCV isolates showed several conserved substitution mutations in specific loci (Supplementary Fig. S2). Mutation of nucleotide acids in virus genome has been well reported to contribute to overcoming host resistance for different viruses ^[44–46]^. These mutations in TYLCV-YN6244 genome likely enable it to break bred-resistance in these tomato cultivars.

**Fig. 1.**
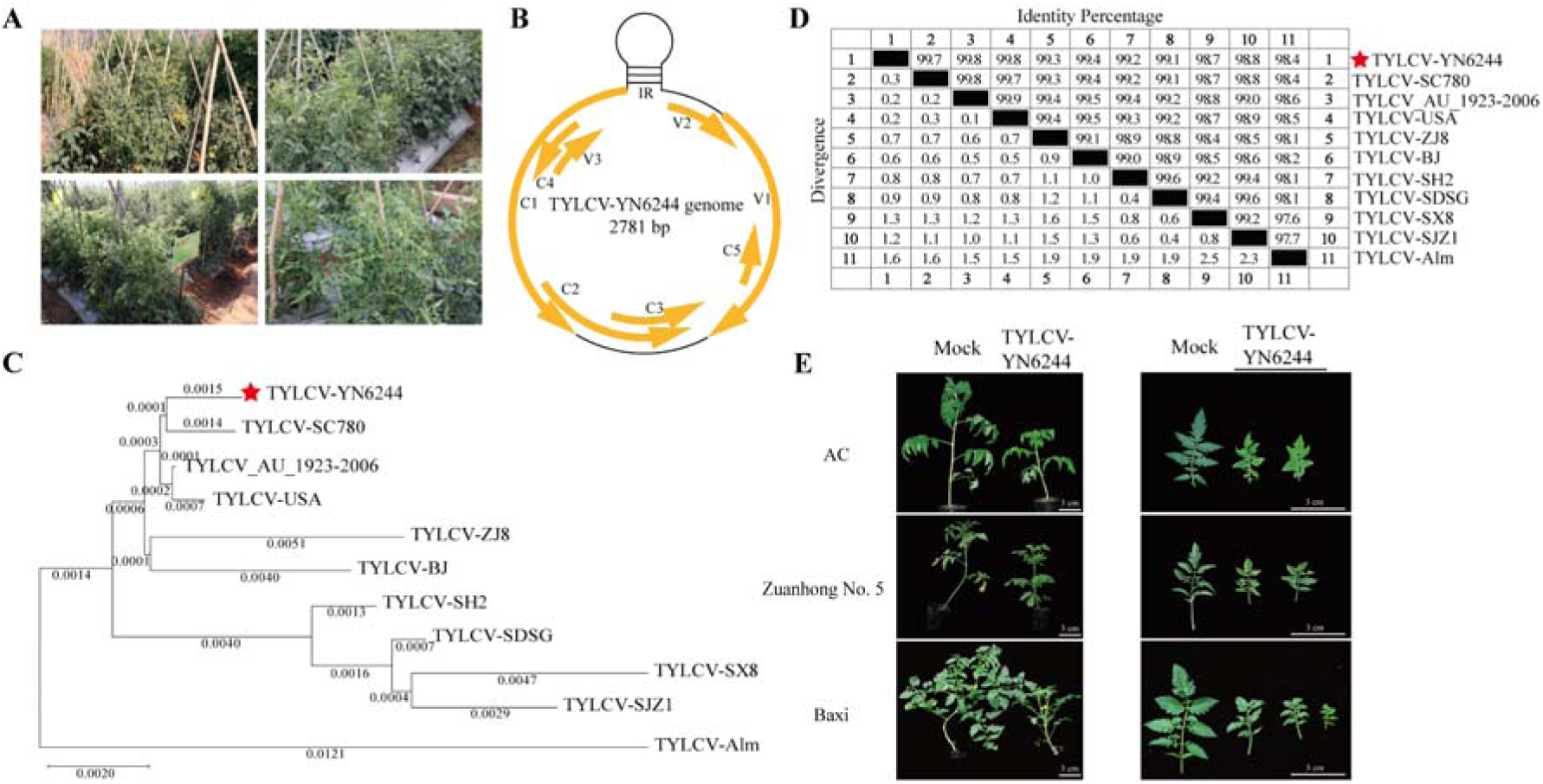
Identification of a novel TYLCV isolate causing disease in resistant tomato cultivars. **A** Disease symptom in TYLCV-YN6244-infected tomato cultivar plants. **B** Schematic diagram of the TYLCV-YN6244 genome. The ORFs (V1, V2, V3 in viral sense and C1, C2, C3, C4, C5 in complementary sense) are indicated with orange curved arrows. **C** Phylogenetic tree analysis of TYLCV-YN6244 with other TYLCV isolates based on the complete genome sequence through MEGA 11. **D** Identity percentage of TYLCV-YN6244 with other TYLCV isolates based on alignment of complete genome sequence using DNASTAR. **E** Phenotype of AC, Zuanhong No.5 and Baxi infected with TYLCV-YN6244 infectious clone. Whole Plants (left panel) and systemic leaves (right panel) were photographed at 50 dpi after Mock treatment or infection with TYLCV-YN6244 infectious clone.

We next conducted a phylogenetic analysis using the complete genome sequences of TYLCV-YN6244 together with other known TYLCV (Supplementary Table S3). It was found that TYLCV-YN6244 showed very close evolutionary relationship with other known TYLCV, especially TYLCV_AU_1923-2006 and TYLCV-USA (Fig. 1C). Further analyses of identity and divergence based on whole genome sequence showed that TYLCV-YN6244 shared very high identity to TYLCV examined, with highest identity ∼99.8% (Fig. 1D). These results indicate that TYLCV-YN6244 is a novel isolate rather than a new species of TYLCV.

To rescue TYLCV-YN6244 and confirm its pathogenicity, we generated infectious clone of TYLCV-YN6244 by cloning the full-length of viral genomic sequence into the binary vector pBinPLUS according to previous report ^[47]^. After being inoculated Zuanhong No.5, Baxi or AC tomato cultivar through agrobacterium-mediated method, TYLCV-YN6244 infectious clone caused severe disease symptoms in all of these tomato cultivar plants (Fig. 1E). These indicated that TYLCV-YN6244 emerged as a novel TYLCV isolate breaking bred-resistance to threaten tomato cultivars.

### V2 protein functions as a potent VSR of TYLCV-YN6244

It has been reported that VSR is the determinant factor inducing viral disease in host plants ^[35, 48]^. To find out the VSR encoded in TYLCV-YN6244 genome, we amplified all different ORFs of TYLCV-YN6244 and respectively cloned them into the binary expression vector pCAMBIA3301 (Fig. 2A). Each viral ORF tagged with a 2×Flag epitope at N-terminus was constructed to be driven under the *ACTIN-2* promoter for expression (Fig. 2A). Each expression vector was co-infiltrated with a Green Fluorescent Protein (GFP) expression vector into 16c transgenic *N. benthamiana* plants which are routinely used for VSR assay ^[36]^. At 5 days post-inoculation (dpi), infiltration of positive control with known vector pGD-VSRs (embedded with Tomato bushy stunt tombusvirus P19, Tobacco etch potyvirus HcPro and Barley yellow mosaic hordeivirus γb) induced intense GFP fluorescence in 16c plants ^[38]^, infiltration of Flag-V2 or Flag-C2 vector respectively led to intense or weak GFP fluorescence while no others resulted to visible GFP fluorescence signals, compared to the negative control with Empty Vector (E.V.) (Fig. 2B). These results indicate that V2 protein function as a potent VSR efficiently suppressing gene silencing in 16c transgenic plants, whereas TYLCV-YN6244 C2 also possesses weak VSR activity.

**Fig. 2.**
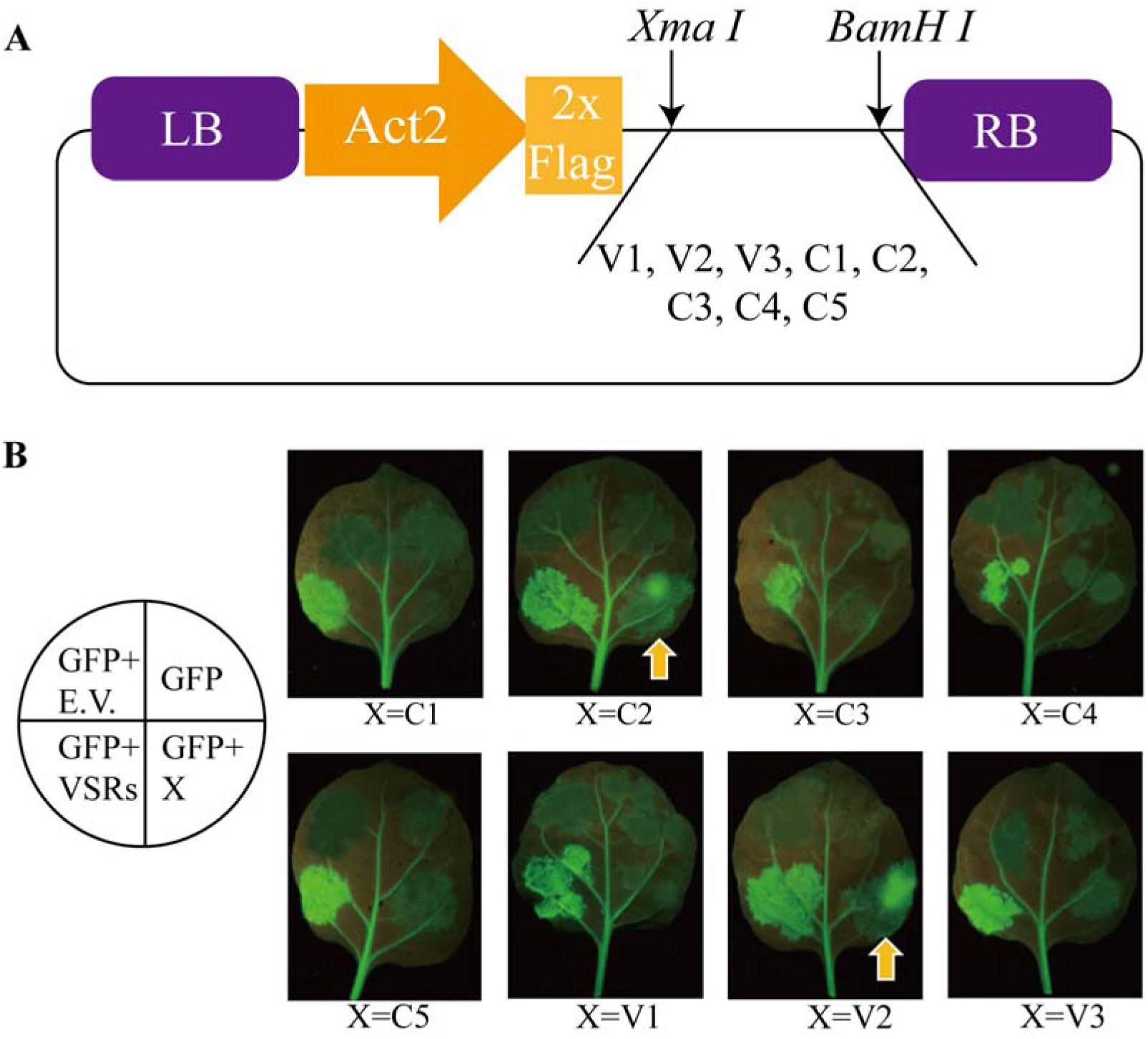
Identification of TYLCV-YN6244 VSR proteins. **A** Schematic diagram for the constructed expression vectors of viral proteins. **B** VSR activity assay in 16c transgenic *N. benthamiana* plants. Each of viral protein vector, together with GFP expression vector, were respectively infiltrated into 16c plants. GFP fluorescence was visualized under ultraviolet light at 5 dpi. Empty vector (E.V.) was used as negative control and vector pGD-VSRs embedded with three different VSRs (P19, HcPro and γb) was used as positive control ^[38]^.

### Infection of TYLCV-YN6244 but not TYLCV-YN6244-**Δ**V2 causes severe disease symptoms and inhibits vsiRNA production in wildtype tobacco plants

We further developed TYLCV-YN6244-ΔV2 infectious clone in which 90 nucleotide acids of *V2* sequence (from 148 to 238) was deleted and overlapping *V1* was intact (Fig. 3A; Supplementary Fig. S3). To assess the infectivity and pathogenicity of TYLCV-YN6244 or TYLCV-YN6244-ΔV2 infectious clone, we then infected them to wild-type *N. benthamiana* plants via Agrobacterium-mediated inoculation. At 21 dpi, plants infected with TYLCV-YN6244 infectious clone exhibited severe disease symptoms, such as leaf curling, chlorosis, and stunted growth, but plants infected with TYLCV-YN6244-ΔV2 infectious clone did not show any visible disease symptom, compared to Mock-treated plants (Fig. 3B). PCR analyses confirmed that both TYLCV-YN6244-ΔV2 and TYLCV-YN6244 systematically infected all *N. benthamiana* plants after inoculation with either infectious clone (Supplementary Table S4). These results not only verified the virulence of TYLCV-YN6244, but also further demonstrated V2 to be a key virulence factor for disease induction in plants.

**Fig. 3.**
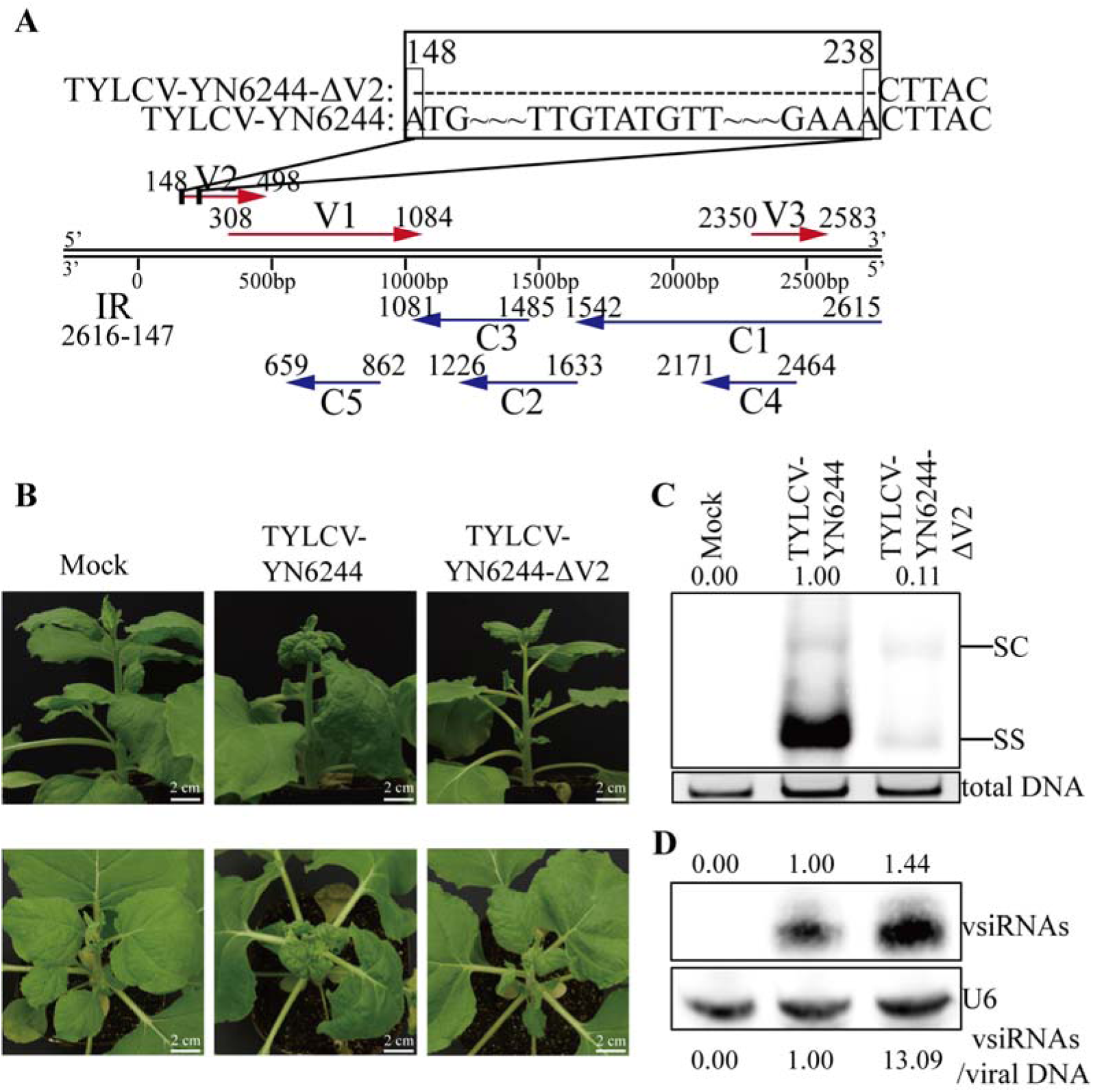
TYLCV-YN6244 but not TYLCV-YN6244-ΔV2 suppresses vsiRNA biogenesis and causes viral disease in tobacco plants. **A** The 90 nt deleted region on *V2* gene for TYLCV-YN6244-ΔV2 was indicated with rectangular box (up), arrows indicate positions of different viral genes in TYLCV-YN6244 genome (down). **B** Phenotype of *N. benthamiana* plants infected with TYLCV-YN6244 or TYLCV-YN6244-ΔV2. Plants were photographed at 21 dpi after Mock treatment or infection with TYLCV-YN6244 or TYLCV-YN6244-ΔV2 infectious clone. **C** Southern blot analyses of viral genomic DNA in tobacco plants at 21 dpi after infection with TYLCV-YN6244 or TYLCV-YN6244-ΔV2. Total DNA was used as the loading control. SC, supercoiled double-stranded DNA; SS, single-stranded DNA. **D** Northern blot analyses of vsiRNA accumulation in tobacco plants at 21 dpi after infection with TYLCV-YN6244 or TYLCV-YN6244-ΔV2. U6 RNA was used as a loading control. Values at the top of the panels represent relative hybridization signal intensities of viral DNA or vsiRNAs, normalized to the loading control. Values at the bottom panel indicate the ratio of vsiRNAs/viral genomic DNA. The signal intensity or ratio in plants infected with TYLCV-YN6244 was set as 1 for relative accumulation calculation. The experiments in (**C**) and (**D**) were independently repeated three times with similar results.

Further, we respectively examined the accumulation of viral genomic DNA and vsiRNAs in these *N. benthamiana* plants. Southern blot analyses showed that viral genomic DNA accumulated drastically more in TYLCV-YN6244-infected plants than those infected with TYLCV-YN6244-ΔV2 (Fig. 3C; Supplementary Fig. S6). In contrast, Northern blot analyses revealed that vsiRNAs were less produced in TYLCV-YN6244-infected plants compared to TYLCV-YN6244-ΔV2-infected plants (Fig. 3D; Supplementary Fig. S7). Further calculation of the ratio of vsiRNAs/viral DNA showed that, the ratio of vsiRNAs/viral DNA was over 10 times higher in TYLCV-YN6244-ΔV2-infected plants than that in TYLCV-YN6244-infected plants. These indicate that V2 potently inhibits vsiRNA biogenesis in plants infected with wildtype TYLCV-YN6244, which probably underlies its roles in virus virulence and pathogenicity in plants.

### Distinct pathogenicity of TYLCV-YN6244 or TYLCV-YN6244-**Δ**V2 and their impact on vsiRNA biogenesis in tomato

We further evaluated the infectivity and pathogenicity of TYLCV-YN6244 and TYLCV-YN6244-ΔV2 in wildtype tomato plants. At 21 dpi, wildtype Micro-Tom plants infected with TYLCV-YN6244 infectious clone exhibited severe disease symptoms, including leaf curling, chlorosis, and stunting, while TYLCV-YN6244-ΔV2-infected plants showed no visible disease symptoms, compared to Mock-treated plants (Fig. 4A). Southern blot analyses showed that viral genomic DNA was accumulated dramatically more in TYLCV-YN6244-infected tomato plants compared to plants infected with TYLCV-YN6244-ΔV2 (Fig. 4B; Supplementary Fig. S8). Notably, real-time quantitative PCR (qPCR) demonstrated that virus accumulation in TYLCV-YN6244-infected plants was over 1000-fold more than that in TYLCV-YN6244-ΔV2-infected plants (Fig. 4D). These results further confirmed the pathogenicity of TYLCV-YN6244 and validated the crucial roles of V2 in virus propagation and virulence in tomato plants.

**Fig. 4.**
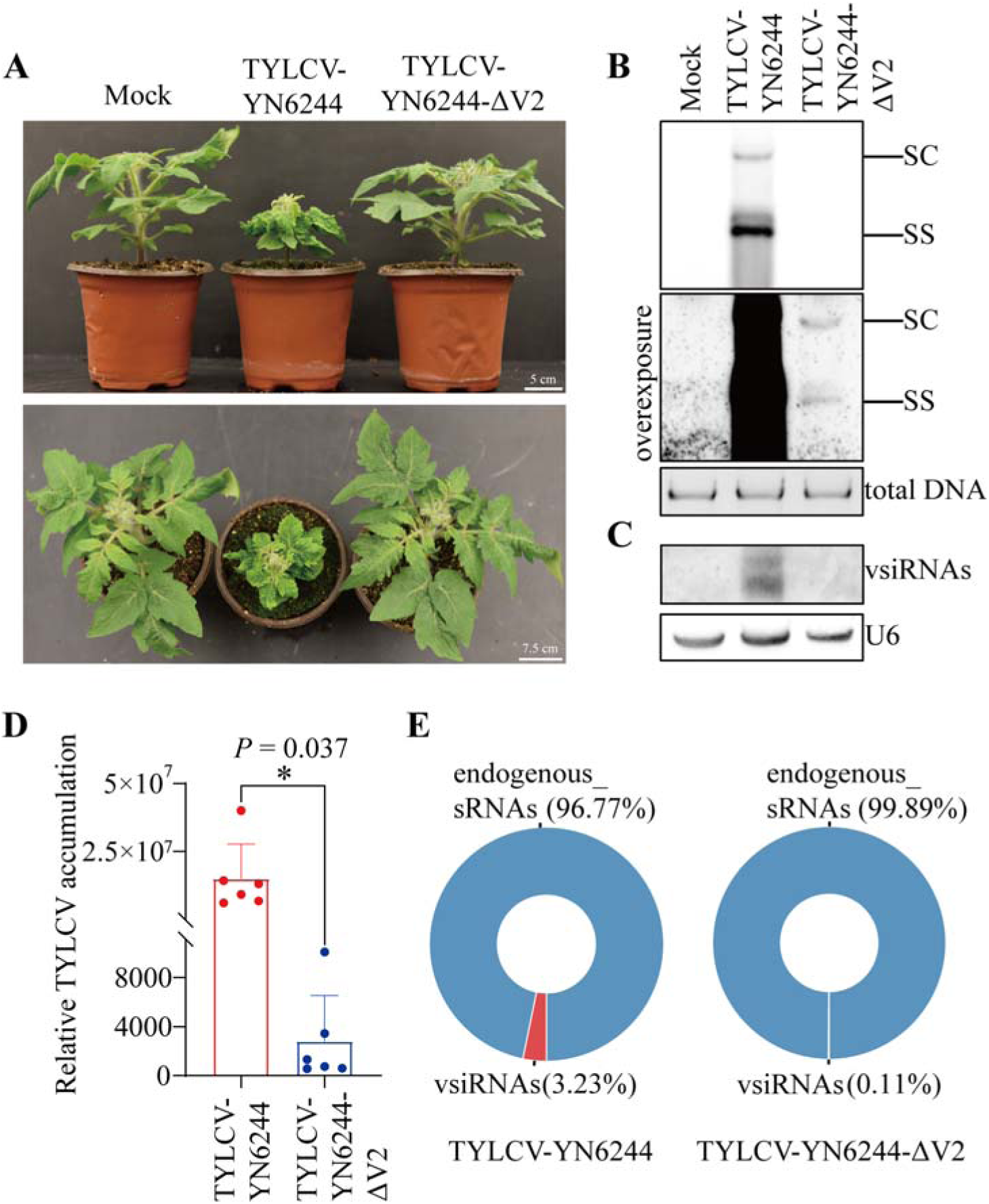
Distinct pathogenicity and impact on vsiRNA biogenesis of TYLCV-YN6244 and TYLCV-YN6244-ΔV2 in tomato. **A** Phenotype of wildtype Micro-tom tomato plants infected with TYLCV-YN6244 or TYLCV-YN6244-ΔV2. Plants were photographed at 21 dpi after Mock treatment, or infection with TYLCV-YN6244 or TYLCV-YN6244-ΔV2 infectious clone. **B** Southern blot analyses of viral genomic DNA in Micro-Tom tomato at 21 dpi after infection with TYLCV-YN6244 or TYLCV-YN6244-ΔV2. Total DNA was used as a loading control. SC, supercoiled double-stranded DNA; SS, single-stranded DNA. **C** Northern blot analyses of vsiRNA accumulation in Micro-Tom tomato at 21 dpi after infection with TYLCV-YN6244 or TYLCV-YN6244-ΔV2. U6 RNA was used as a loading control. **D** qPCR quantification of relative viral DNA accumulation in tomato plants infected with TYLCV-YN6244 or TYLCV-YN6244-ΔV2. Primer pairs for qRT-PCR analyses are listed in Supplementary Table S5. Error bars represent the standard deviation. Asterisks indicate statistically significant differences (*P* < 0.05). **E** Percentages of vsiRNAs or endogenous small RNAs in tomato plants infected with TYLCV-YN6244 or TYLCV-YN6244-ΔV2. The experiments in (**B**) and (**C**) were independently repeated three times with similar results.

Intriguingly, Northern blot analyses showed that vsiRNAs were clearly detected in tomato plants infected with TYLCV-YN6244 but not TYLCV-YN6244-ΔV2 (Fig. 4C; Supplementary Fig. S9), which is different from above finding that vsiRNAs were more efficiently produced in tobacco plants infected with TYLCV-YN6244-ΔV2 (Fig. 3D; Supplementary Fig. S7). We speculated that the low production of vsiRNAs in TYLCV-YN6244-ΔV2-infected plants was probably due to the extremely low accumulation of virus in these plants (Fig. 4B and D; Supplementary Fig. S8). To clarify this, we conducted small RNA-seq analysis on tomato plants after infection with TYLCV-YN6244 or TYLCV-YN6244-ΔV2. It was found that vsiRNA reads were about 3.23% or 0.11% of the total small RNA reads in TYLCV-YN6244-infected plants or TYLCV-YN6244-ΔV2-infected plants, respectively (Fig. 4E). This indicated that approximately 30 times less of vsiRNAs were produced in plants infected with TYLCV-YN6244-ΔV2 than that in plants infected with TYLCV-YN6244. Since virus accumulation in TYLCV-YN6244-ΔV2-infected plants was over 1000 times less than that in TYLCV-YN6244-infected plants (Fig. 4D), the relative efficiency of vsiRNA biogenesis in TYLCV-YN6244-ΔV2-infected plants was conversely increased by dozens of times compared to that in TYLCV-YN6244-infected plants. These results show that TYLCV-YN6244 V2 also functions to suppress vsiRNA biogenesis in tomato.

### 21nt and 22nt rather than 24nt vsiRNAs targeting specific viral genes were predominantly produced in tomato after TYLCV-YN6244 or TYLCV-YN6244-**Δ**V2 infection

Interestingly, profiling vsiRNAs revealed that the predominant 21nt and 22nt rather than 24nt vsiRNAs were produced in tomato plants infected with either TYLCV-YN6244 or TYLCV-YN6244-ΔV2 (Fig. 5A). In TYLCV-YN6244-infected plants, about 46.16% and 24.86% of total vsiRNAs were respectively 21nt and 22nt vsiRNAs, while 24nt vsiRNAs were only accounted for about 6.86% of total vsiRNAs (Fig. 5B). And in TYLCV-YN6244-ΔV2-infected plants, 49.69% and 32.36% of total vsiRNAs were respectively 21nt and 22nt vsiRNAs, while 24nt vsiRNAs were only about 5.62% (Fig. 5B).

**Fig. 5.**
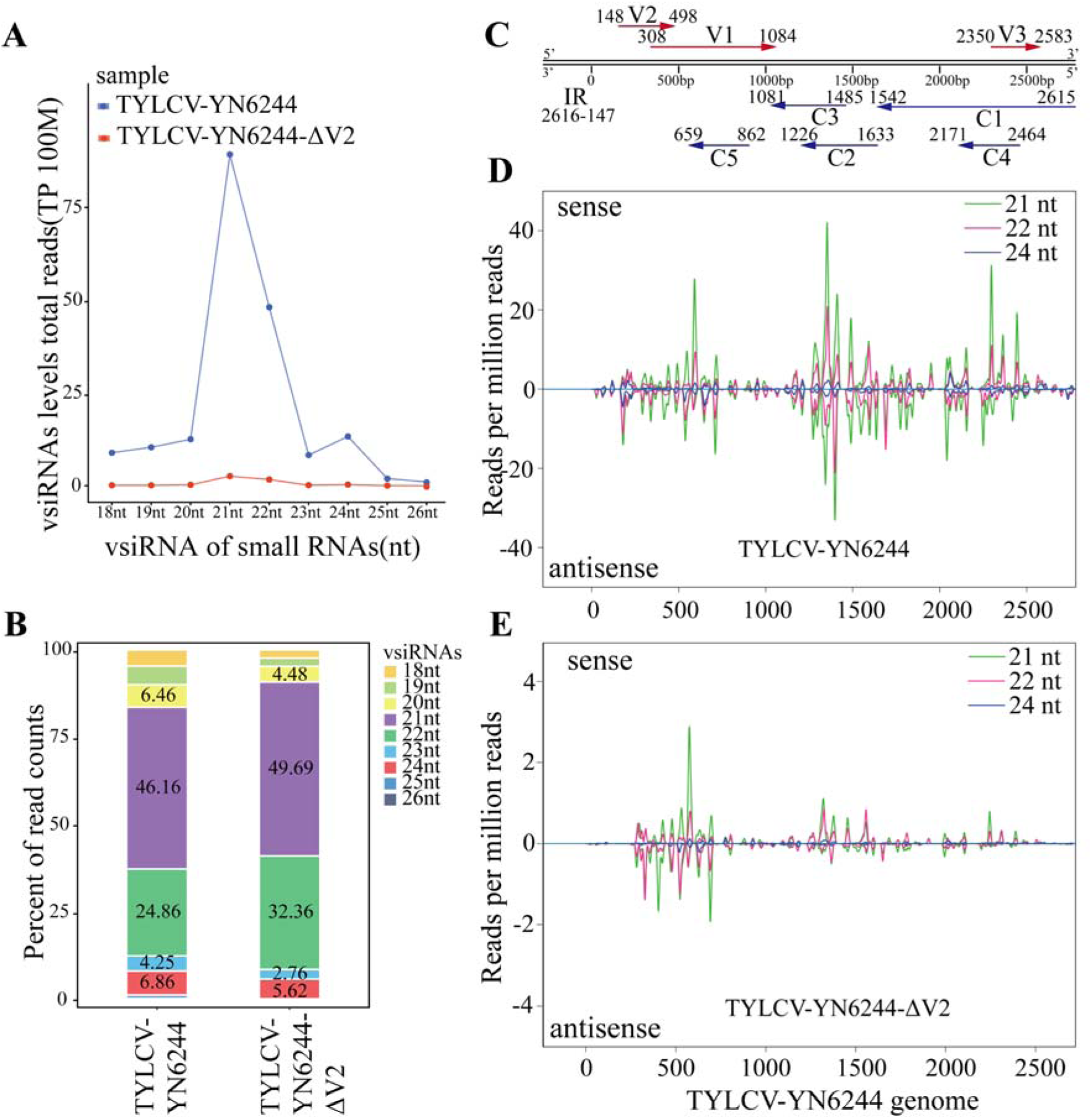
vsiRNA profiling of tomato plants infected with TYLCV-YN6244 or TYLCV-YN6244-ΔV2. **A** Relative quantity of different length-size vsiRNAs in tomato based on small RNA-seq analyses. nt = nucleotides; TPM = transcripts per million. **B** Percentages of different length-size vsiRNAs in tomato plants infected with TYLCV-YN6244 or TYLCV-YN6244-ΔV2. **C** Diagram of genome structure of TYLCV-YN6244. ORFs and their orientations are indicated with arrowed lines. Numbers on ORFs indicate the start and termination site of each ORF. **D, E** Distribution mapping of 21nt, 22nt, and 24nt vsiRNAs on the TYLCV-YN6244 (**D**) or TYLCV-YN6244-ΔV2 (**E**) genome.

Furthermore, it was found that 21nt and 22nt vsiRNAs could be mapped to the whole TYLCV-YN6244 or TYLCV-YN6244-ΔV2 genome, particularly the region of viral coding genes such as *V1* and *C3* (Fig. 5C-E). These findings were different from published reports that 24nt vsiRNAs were abundantly produced after TYLCV infection in plants and mainly targeted Intergenic region (IR) of TYLCV ^[49, 50]^. It is very likely that 21nt and 22nt vsiRNAs play distinct roles in small RNA-mediated antiviral defense against plant DNA viruses in tomato.

### Differential modulation of metabolic and defense pathways in tomato by TYLCV-YN6244 and TYLCV-YN6244-**Δ**V2

To further explore the molecular basis of the differences in viral pathogenicity and disease symptoms induced in plants, we performed RNA-seq to analyze transcriptome in tomato plants infected with TYLCV-YN6244 or TYLCV-YN6244-ΔV2. Compared to Mock plants, a total of 1292 and 948 differentially expressed genes (DEGs) were respectively up-regulated and down-regulated in TYLCV-YN6244-infected plants (Supplementary Fig. S4A and B), and 1357 and 834 DEGs were respectively up-regulated and down-regulated in TYLCV-YN6244-ΔV2-infected plants (Supplementary Fig S4C and D). Venn analysis showed that 696 DEGs were concurrently up-regulated in TYLCV-YN6244 or TYLCV-YN6244-ΔV2 infected plants, while 596 and 661 DEGs were uniquely up-regulated in TYLCV-YN6244 or TYLCV-YN6244-ΔV2 infected plants, respectively (Fig. 6A). On the other hand, 441 DEGs were concurrently down-regulated in tomato plants after the infection of TYLCV-YN6244 or TYLCV-YN6244-ΔV2, but 507 or 393 DEGs were respectively down-regulated after TYLCV-YN6244 or TYLCV-YN6244-ΔV2 infection (Fig. 6B). These indicated that transcriptomes in tomato plants were dramatically modulated after the infection of either TYLCV-YN6244 or TYLCV-YN6244-ΔV2.

**Fig. 6.**
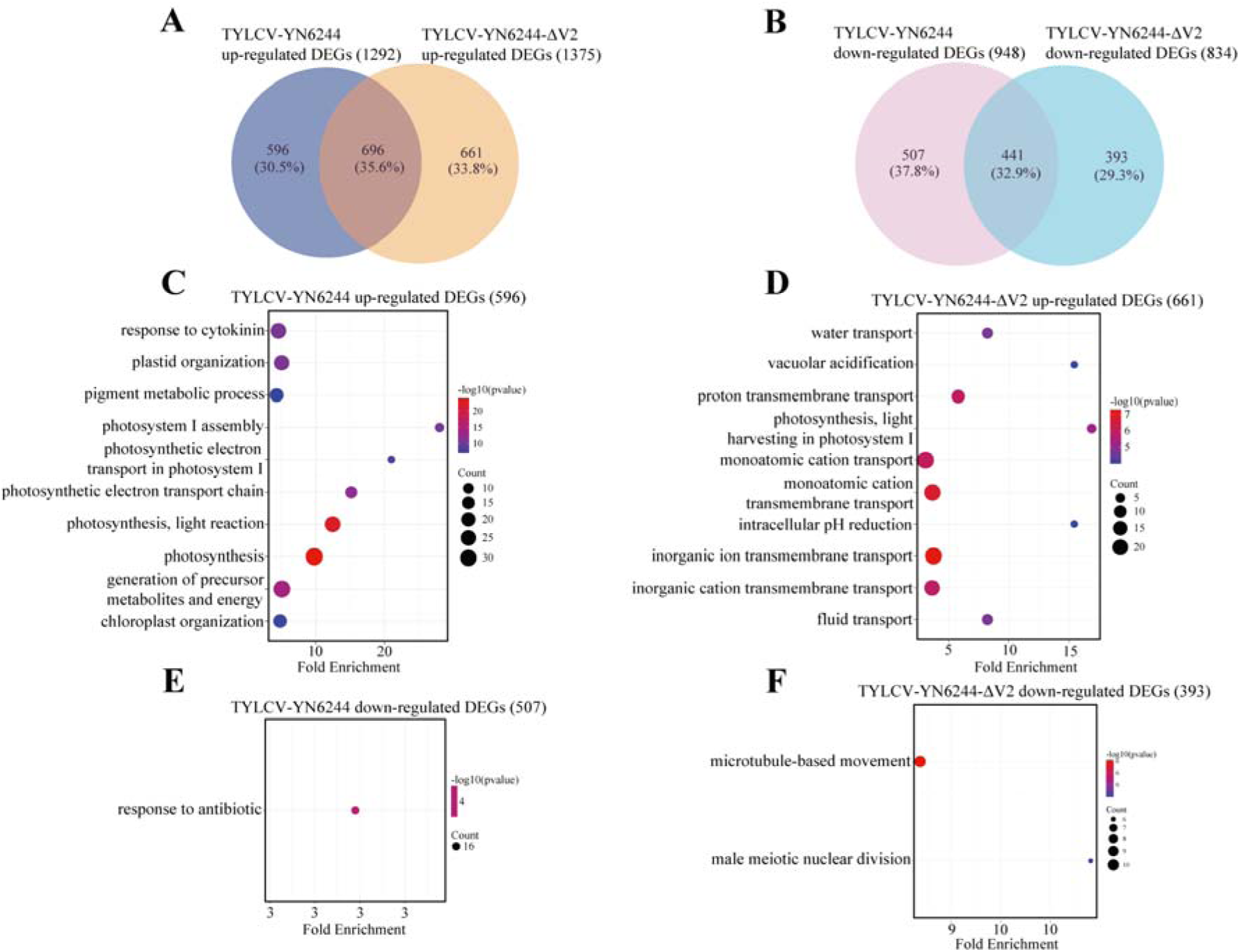
Differential modulation of transcriptome in tomato infected with TYLCV-YN6244 or TYLCV-YN6244-ΔV2. **A** Venn analysis of upregulated DEGs after TYLCV-YN6244 or TYLCV-YN6244-ΔV2 infection. **B** Venn analysis of downregulated DEGs after TYLCV-YN6244 or TYLCV-YN6244-ΔV2 infection. **C - F** GO enrichment analysis of specifically upregulated DEGs (**C** and **D**) or downregulated DEGs (**E** and **F**) in tomato infected with TYLCV-YN6244 or TYLCV-YN6244-ΔV2.

Further, Gene Ontology (GO) analyses were conducted to find out whether specific pathways were modulated by either virus. It was found that, upon the infection of both viruses, the concurrently upregulated DEGs were enriched by genes related to oxidative stress and photosynthesis (Supplementary Fig S5A). However, the exclusively upregulated DEGs after TYLCV-YN6244 infection was associated with cytokinin response, pigment metabolism, and photosynthesis, while genes related to water and ion transport were specifically upregulated after TYLCV-YN6244-ΔV2 infection (Fig. 6C and D), which is consistently with the severe disease symptom after TYLCV-YN6244 infection. On the other hand, the concurrently downregulated DEGs were enriched by genes involved in floral organ development and meristem determinacy (Supplementary Fig. S5B). But, the exclusively downregulated DEGs after TYLCV-YN6244 infection was associated with antibiotic response, while genes related to microtubule-based movement and male meiotic nuclear division were specifically downregulated after TYLCV-YN6244-ΔV2 infection (Fig. 6E and F). These results indicated that various development or defense pathways were differentially modulated in tomato after the infection of TYLCV-YN6244 or TYLCV-YN6244-ΔV2, which together probably contributed to the distinct disease symptoms and viral accumulation in tomato plants after the infection of either virus.

## Discussion

TYLCD, caused by different species of TYLCV, remains to be one of the most devastating viral diseases worldwide ^[51]^. Although resistant tomato cultivars have been bred to prevent TYLCD, viruses evolve rapidly to overcome host resistance and adapt to diverse environmental conditions ^[52]^. In this study, we identified a novel TYLCV isolate-TYLCV-YN6244 at Yunnan province of China, causing severe disease in resistant tomato cultivars (Fig. 1). TYLCV-YN6244 could pose a significant threat to tomato production and request to develop new resistance in germplasms.

We determined the complete nucleotide sequence of TYLCV-YN6244, which showed a very high homology with other TYLCV isolates, and encoded six typical viral proteins of Begomoviruses (Fig. 1). Further investigation identified V2 as a potent VSR with strong suppression activities (Fig. 2). Further, we generated infectious clones of wildtype TYLCV-YN6244 and VSR V2-deficient infectious clone (TYLCV-YN6244-ΔV2) (Figs. 3A). We demonstrated that, both TYLCV-YN6244 and TYLCV-YN6244-ΔV2 high efficiently and systemically infected tobacco and tomato plants, however, TYLCV-YN6244-ΔV2 failed to induce disease symptoms in tobacco or tomato plants, with dramatically reduced viral accumulation, compared to TYLCV-YN6244 (Fig. 3C and 4D). These results verified virulence and pathogenicity with developed infectious clone and demonstrated critical roles of V2 in viral virulence and pathogenicity in plants, probably by suppressing antiviral RNAi through inhibiting vsiRNA biogenesis.

vsiRNAs are the product of the antiviral defense process that is naturally invoked by a host when perceiving viral invasion ^[24]^. Plants usually produce vsiRNAs through Dicer-like proteins (DCLs). In *Arabidopsis thaliana*, viral RNAs are cleaved primarily by DCL4 and secondarily by DCL2, which results in 21- and 22-nucleotide (nt) vsiRNAs, respectively, which are loaded into AGO1 or AGO2 to mediate RNA silencing ^[53, 54]^. In addition, the 24-nucleotide vsiRNAs are mainly produced by DCL3 and play a key role in RNAi-mediated plant DNA viral defense through the TGS pathway ^[55, 56]^. In tomato, however, we found that 21nt and 22nt vsiRNAs, but not 24nt vsiRNAs, were predominantly produced after TYLCV-YN6244 or TYLCV-YN6244-ΔV2 infection (Fig. 5B) and could be mapped to TYLCV-YN6244 whole genome, particularly on *V1* and *C3* regions (Fig. 5C). It is worth to note that the 21 and 22nt vsiRNAs have also been identified upon the infection of plant DNA viruses in tomato ^[54]^. These suggested that 21nt and 22nt vsiRNAs may also play vital roles in the defense of plant DNA viruses including TYLCV in tomato. It is well known that both 21nt and 22nt vsiRNAs are primarily involved in PTGS-mediated antiviral defense against RNA viruses ^[23]^, their roles in DNA virus defense have not been well elucidated. 21nt vsiRNAs have been reported to be implicated in RNA-dependent DNA methylation (RdDM) which is implicated in antiviral defense to plant DNA viruses through TGS pathway in plants ^[29]^. However, it is also possible that these 21nt and 22nt vsiRNAs mediate slicing or translation inhibition of viral RNA through PTGS pathway in defense of plant DNA viruses. Tomato contains multiple homologs of DCL2, AGO1, AGO2, and one homolog of DCL4 ^[57]^. Characterizing the function of these components of canonical PTGS pathway will contribute to clarifying 21nt and 22nt vsiRNAs-mediated antiviral defense to plant DNA viruses in the future.

Notably, a robust genetic screen using VSR-deficient CMV (CMV-Δ2b) has been established recently to elucidate the antiviral defense against plant RNA viruses ^[26, 27, 32, 58]^. However, a comparable approach for studying antiviral RNAi against plant DNA viruses is still missing. In this study, a novel TYLCV isolate (TYLCV-YN6244) of plant DNA virus with significant agricultural threat was identified and characterized. We generated not only an infectious clone of wildtype TYLCV-YN6244, but also a VSR V2-deficient infectious clone of TYLCV (TYLCV-YN6244-ΔV2) in which *V2* was completely but solely dysfunctional (Fig. 3A; Supplementary Fig. S3). These will provide important resources for developing a comparable genetic system to dissect antiviral RNAi against plant DNA viruses in *Solanaceae* plants, eventually contributing to breeding new resistance in agricultural crops.

Together, our research here not only identified and characterized a novel pathogenic TYLCV isolate overcoming bred-resistance to threaten tomato cultivar and but also developed important resources to study the interaction of virus and host plants for breeding novel resistance in important crop plants.

## List of abbreviations

TYLCV: Tomato yellow leaf curl virus
VSR: Viral suppressor of RNA silencing
vsiRNAs: Virus-derived small interfering RNAs
ORF: Open reading frames
IR: Intergenic region
CP: Coat protein
MP: Movement protein
PTGS: Post-transcriptional gene silencing
TGS: Transcriptional gene silencing
TYLCD: Tomato yellow leaf curl disease
CMV: Cucumber mosaic virus
TSWV: Tomato spotted wilt virus
DCL: Dicer-like
ds-RNA: Double-stranded RNA
AVI: Antiviral RNAi-defective
AGO: Argonaute
RISC: RNA-induced silencing complex
RDR: RNA-dependent RNA polymerases
RdDM: RNA-directed DNA methylation
Dpi: Days post-inoculation
qPCR: Real-time quantitative PCR
RNA-seq: RNA sequencing
sRNA-seq: Small RNA sequencing
DEGs: Differentially expressed genes
GO: Gene Ontology
ROS: Reactive oxygen species

## Supplementary Information

**Additional file 1: Supplementary Fig. S1** Identification of *Ty-1* in Zuanhong No. 5 (**A**) and *Ty-2* in Baxi (**B**). M represent Marker, N represent Native control, P represent Positive control. Primer pairs are listed in Supplementary Table S5. **Fig. S2** Multiple alignments of TYLCV-YN6244 with other TYLCV isolates. Mutation sites in TYLCV-YN6244 genome are indicated by red rectangular boxes. **Fig. S3** Procedure diagram of constructing the infectious clones of TYLCV-YN6244 and TYLCV-YN6244-ΔV2. 0.5-unit and 1.0-unit tandem repeats of TYLCV-YN6244 were cloned into the plant binary vector pBinPLUS. The restriction enzymes used for cloning were highlighted in red. **Fig. S4** Differentially expressed genes (DEG) in tomato infected with TYLCV-YN6244 or TYLCV-YN6244-ΔV2. **A, C** Volcano plots representing DEGs in tomato infected with TYLCV-YN6244 (**A**) or TYLCV-YN6244-ΔV2 (**C**). **B, D** Expression pattern clustering heatmap of DEGs in tomato infected with TYLCV-YN6244 (**B**) or TYLCV-YN6244-ΔV2 (**D**). **Fig. S5** GO enrichment of concurrently up-regulated DEGs (**A**) or concurrently down-regulated DEGs (**B**) in tomato inoculated with TYLCV-YN6244 or TYLCV-YN6244-ΔV2. **Fig. S6** TYLCV-YN6244 but not TYLCV-YN6244-ΔV2 suppresses vsiRNA biogenesis and causes viral disease in tobacco plants, related to Fig. 3. The red dashed box corresponding to Fig. 3C. Southern blot analyses of viral genomic DNA in tobacco plants at 21 dpi after infection with TYLCV-YN6244 or TYLCV-YN6244-ΔV2. Total DNA was used as the loading control. SC, supercoiled double-stranded DNA; SS, single-stranded DNA. **Fig. S7** TYLCV-YN6244 but not TYLCV-YN6244-ΔV2 suppresses vsiRNA biogenesis and causes viral disease in tobacco plants, related to Fig. 3. The red dashed box corresponding to Fig. 3D. Northern blot analyses of vsiRNA accumulation in tobacco plants at 21 dpi after infection with TYLCV-YN6244 or TYLCV-YN6244-ΔV2. U6 RNA was used as a loading control. **Fig. S8** Distinct pathogenicity and impact on vsiRNA biogenesis of TYLCV-YN6244 and TYLCV-YN6244-ΔV2 in tomato, related to Fig. 4. The red dashed box corresponding to Fig. 4B. Southern blot analyses of viral genomic DNA in Micro-Tom tomato at 21 dpi after infection with TYLCV-YN6244 or TYLCV-YN6244-ΔV2. Total DNA was used as a loading control. SC, supercoiled double-stranded DNA; SS, single-stranded DNA. **Fig. S9** Distinct pathogenicity and impact on vsiRNA biogenesis of TYLCV-YN6244 and TYLCV-YN6244-ΔV2 in tomato, related to Fig. 4. The red dashed box corresponding to Fig. 4C. Northern blot analyses of vsiRNA accumulation in Micro-Tom tomato at 21 dpi after infection with TYLCV-YN6244 or TYLCV-YN6244-ΔV2. U6 RNA was used as a loading control.

**Additional file 2: Supplementary Table S1** TYLCV-YN6244 genome sequence. **Table S2** Coding genes and Intergenic region (IR) of TYLCV-YN6244. **Table S3** Known TYLCV isolates used for phylogenetic tree analysis. **Table S4** PCR analyses of systemic infection rate of TYLCV-YN6244 and TYLCV-YN6244-ΔV2 in tobacco plants. PCR analysis was conducted via primer pairs of TYLCV-YN6244-IR-F/R. **Table S5** Primers used in this study.

## Acknowledgements

We thank professor Xianbing Wang (College of Biological Sciences, China Agricultural University, China) for providing the pGD-VSRs and all members in our Lab for helpful suggestion and support.

## Authors’ contributions

ZXG and LLZ conceived the study; YGZ, SHJ, XMX and JZ performed the experiments; YGZ and SHJ performed data analysis, YGZ wrote the original manuscript, ZXG, LLZ and MD supervised the experiments and revised the manuscript. All authors read and approved the final manuscript.

## Availability of data

The raw sequence data reported in this paper have been deposited in the Genome Sequence Archive (Genomics, Proteomics & Bioinformatics 2025) in National Genomics Data Center (Nucleic Acids Res 2025), China National Center for Bioinformation / Beijing Institute of Genomics, Chinese Academy of Sciences (GSA: CRA037679 and CRA037680) that are publicly accessible at https://ngdc.cncb.ac.cn/gsa.

## Declarations

### Ethics approval and consent to participate

Not applicable.

### Consent for publication

Not applicable.

