## Supplementary Fig for "Identification and molecular characterization of a novel TYLCV isolate breaking bred-resistance to threaten tomato cultivar"

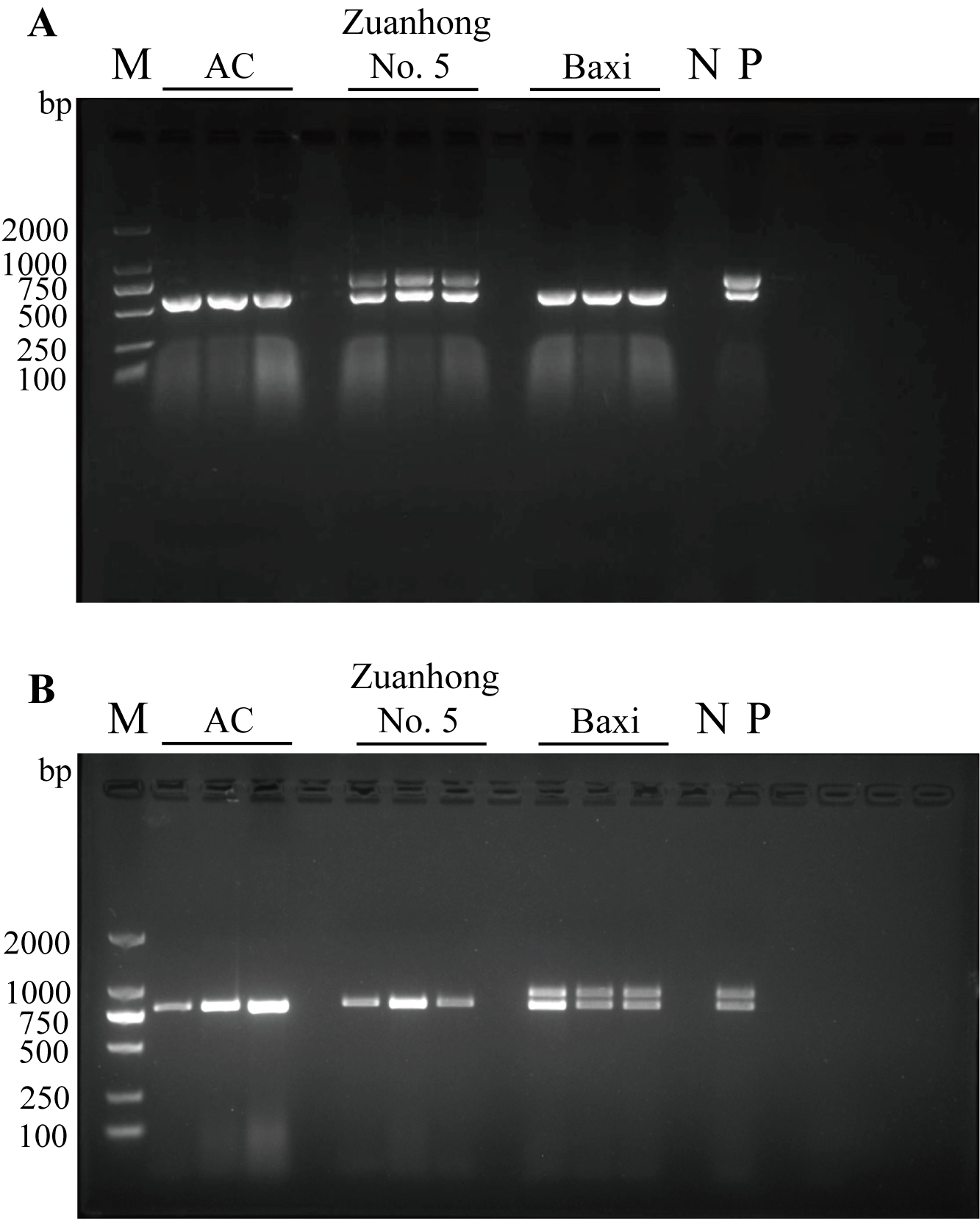
**Fig. S1**	Identification of *Ty-1* in Zuanhong No. 5 (**A**) and *Ty-2* in Baxi (**B**). M represent Marker, N represent Native control, P represent Positive control. Primer pairs are listed in Supplementary Table S5.

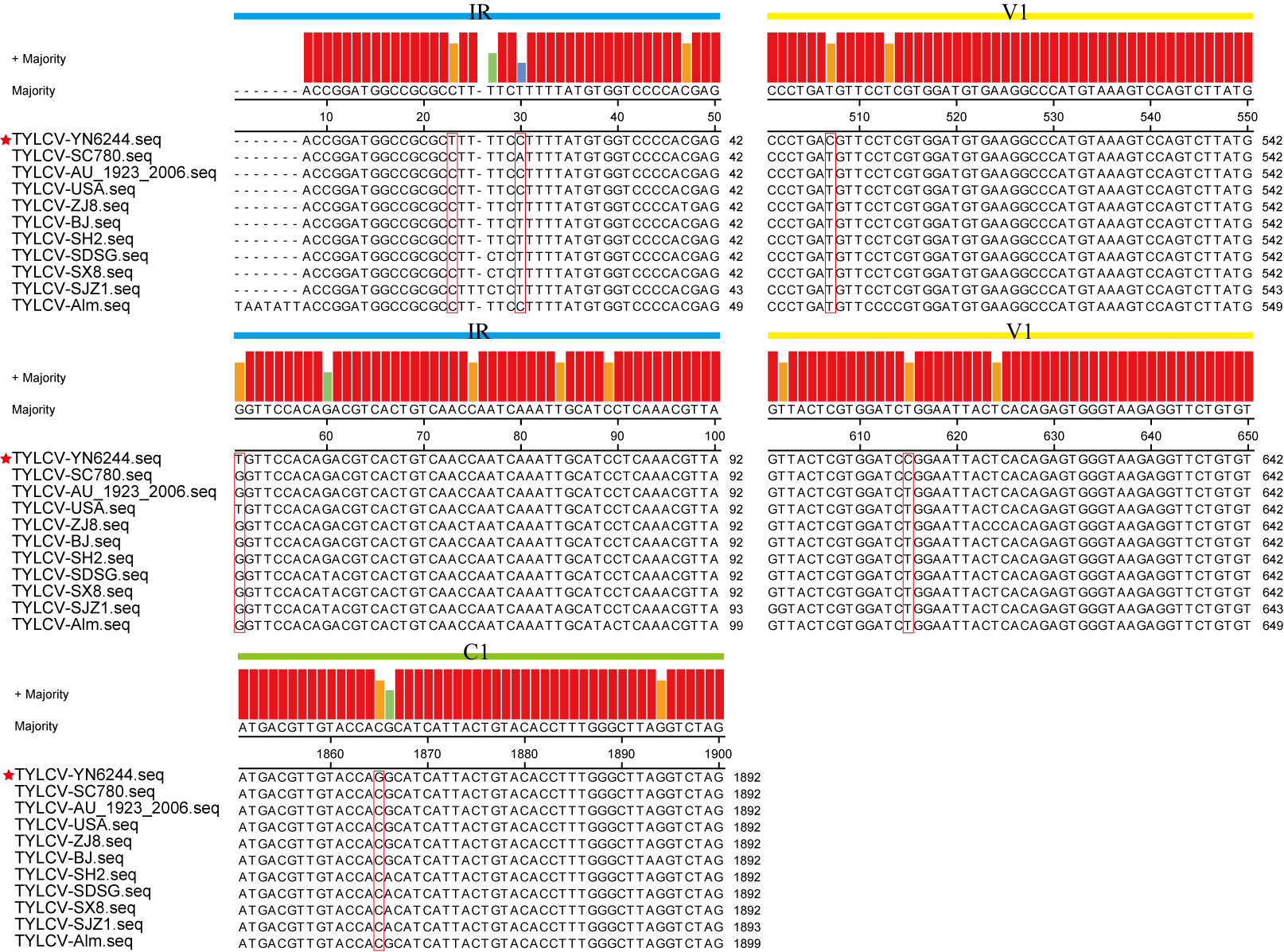

**Fig. S2** Multiple alignments of TYLCV-YN6244 with other TYLCV isolates. Mutation sites in TYLCV-YN6244 genome are indicated by red rectangular boxes.

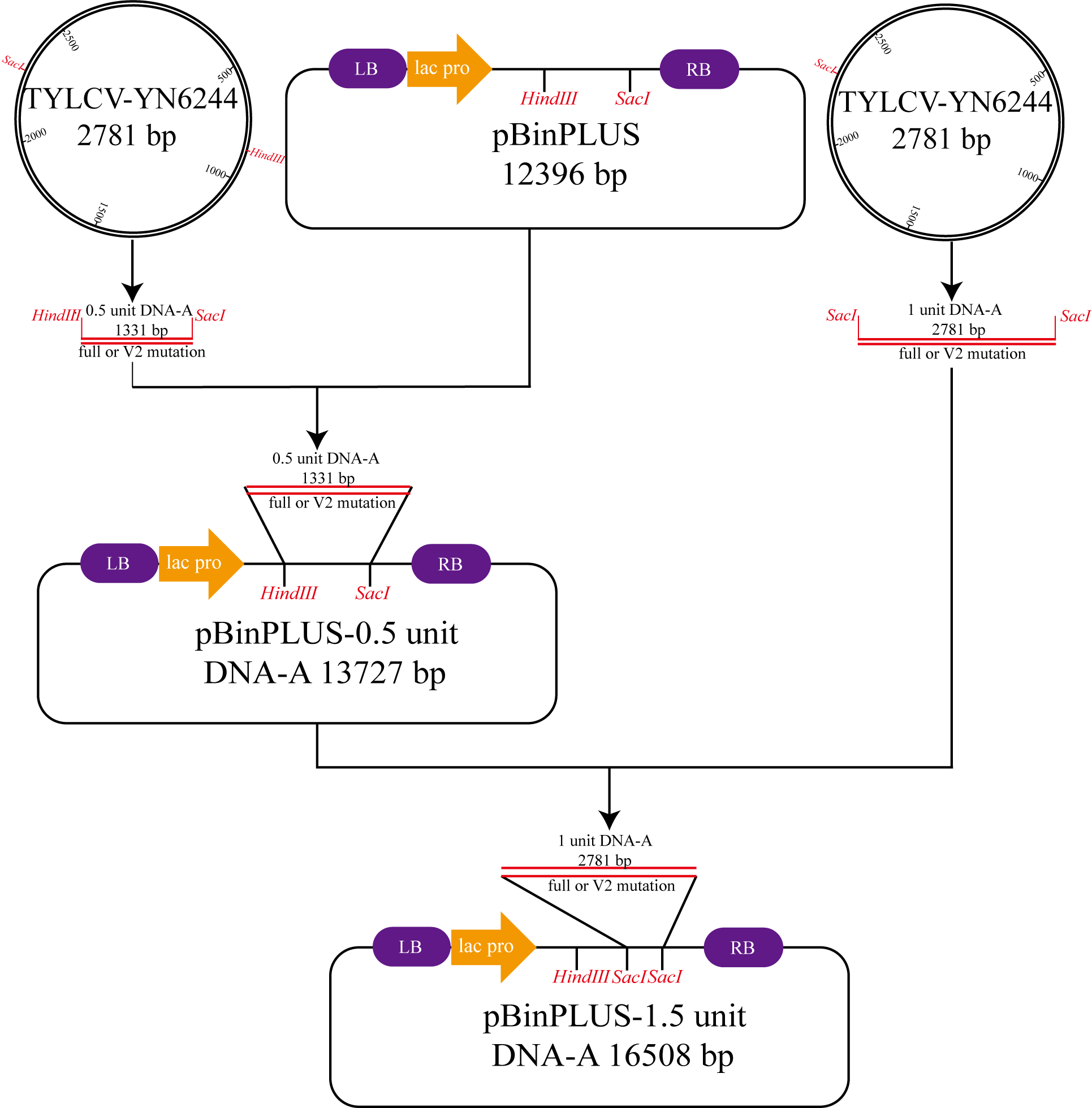

**Fig. S3** Procedure diagram of constructing the infectious clones of TYLCV-YN6244 and TYLCV-YN6244-ΔV2. 0.5-unit and 1.0-unit tandem repeats of TYLCV-YN6244 were cloned into the plant binary vector pBinPLUS. The restriction enzymes used for cloning were highlighted in red.

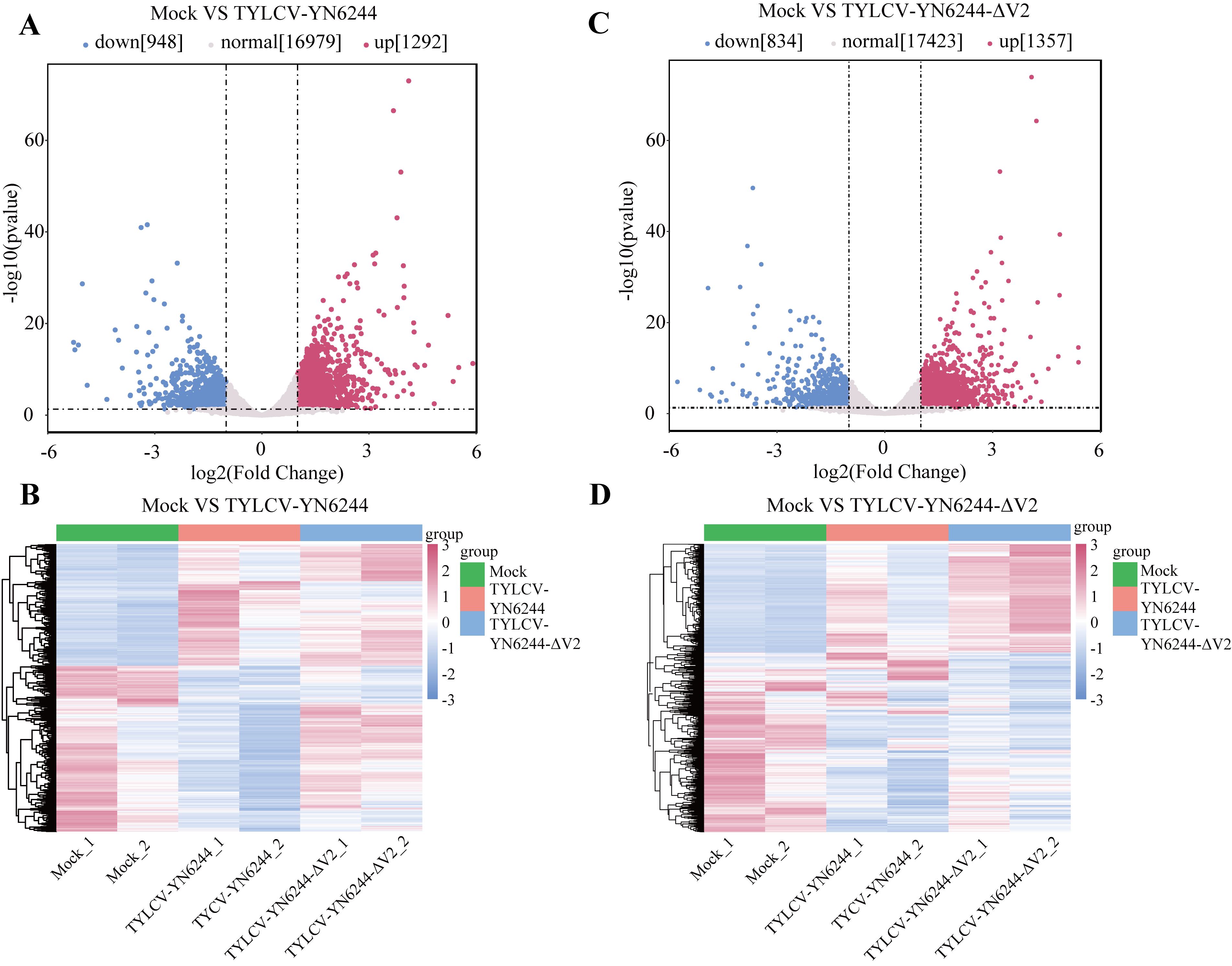

**Fig. S4** Differentially expressed genes (DEG) in tomato infected with TYLCV-YN6244 or TYLCV-YN6244-ΔV2. **A, C** Volcano plots representing DEGs in tomato infected with TYLCV-YN6244 (**A**) or TYLCV-YN6244-ΔV2 (**C**). **B, D** Expression pattern clustering heatmap of DEGs in tomato infected with TYLCV-YN6244 (**B**) or TYLCV-YN6244-ΔV2 (**D**).

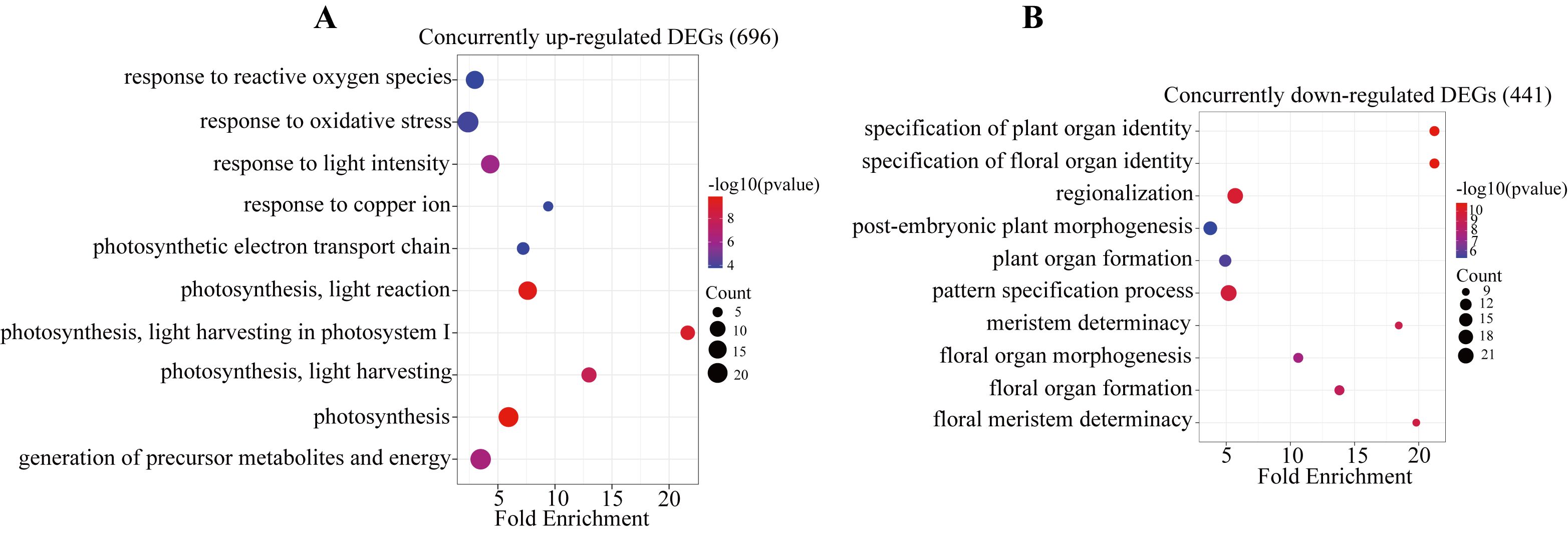

**Fig. S5** GO enrichment of concurrently up-regulated DEGs (**A**) or concurrently down-regulated DEGs (**B**) in tomato inoculated with TYLCV-YN6244 or TYLCV-YN6244-ΔV2.

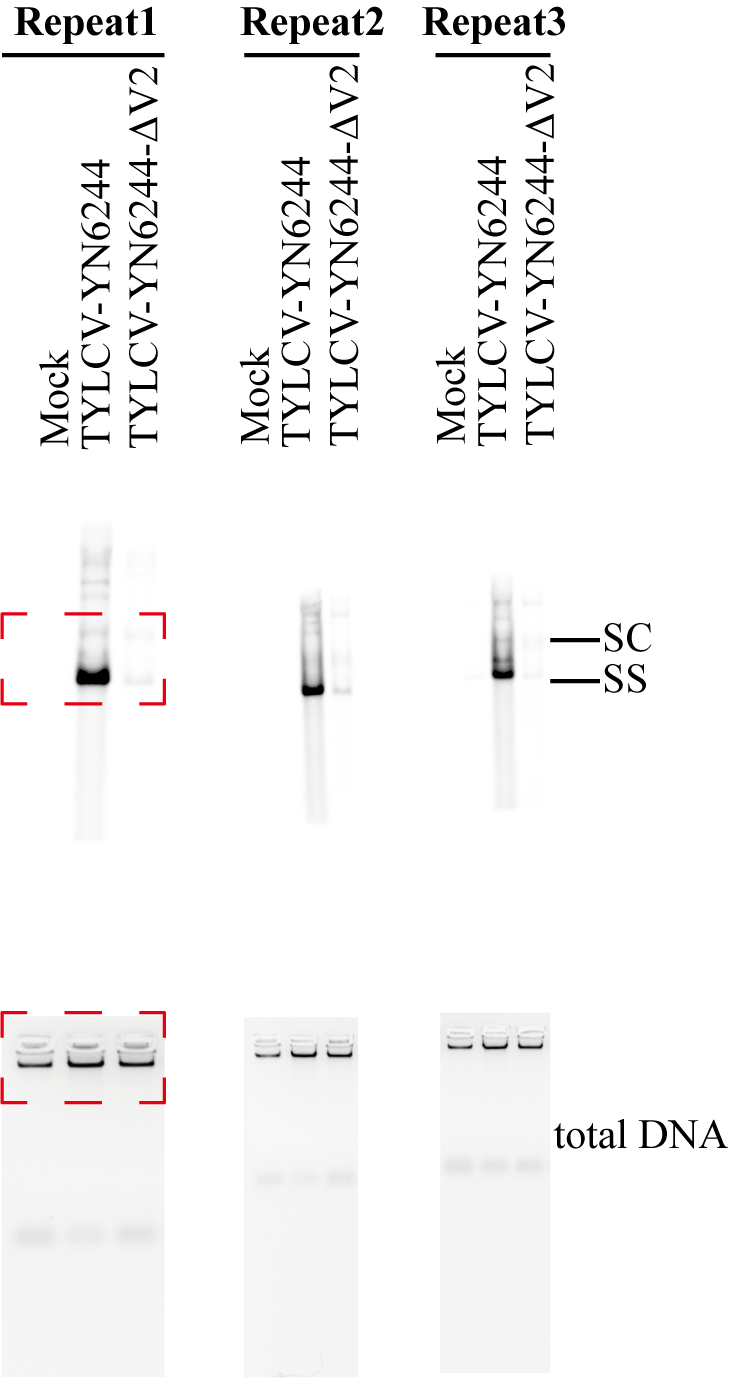
**Fig. S6** TYLCV-YN6244 but not TYLCV-YN6244-ΔV2 suppresses vsiRNA biogenesis and causes viral disease in tobacco plants, related to Fig. 3. The red dashed box corresponding to Fig. 3C. Southern blot analyses of viral genomic DNA in tobacco plants at 21 dpi after infection with TYLCV-YN6244 or TYLCV-YN6244-ΔV2. Total DNA was used as the loading control. SC, supercoiled double-stranded DNA; SS, single-stranded DNA.

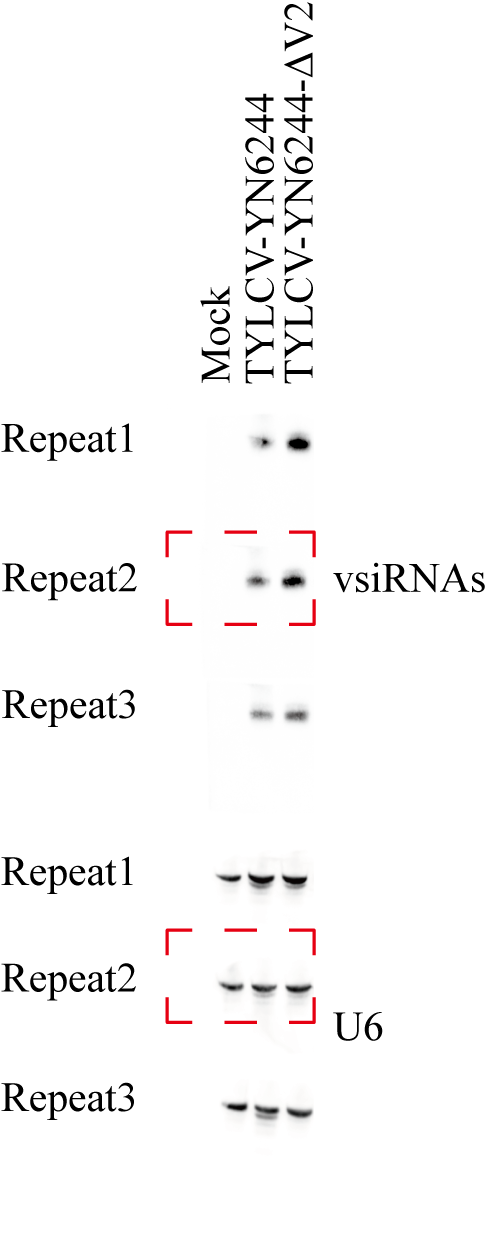
**Fig. S7** TYLCV-YN6244 but not TYLCV-YN6244-ΔV2 suppresses vsiRNA biogenesis and causes viral disease in tobacco plants, related to Fig. 3. The red dashed box corresponding to Fig. 3D. Northern blot analyses of vsiRNA accumulation in tobacco plants at 21 dpi after infection with TYLCV-YN6244 or TYLCV-YN6244-ΔV2. U6 RNA was used as a loading control.

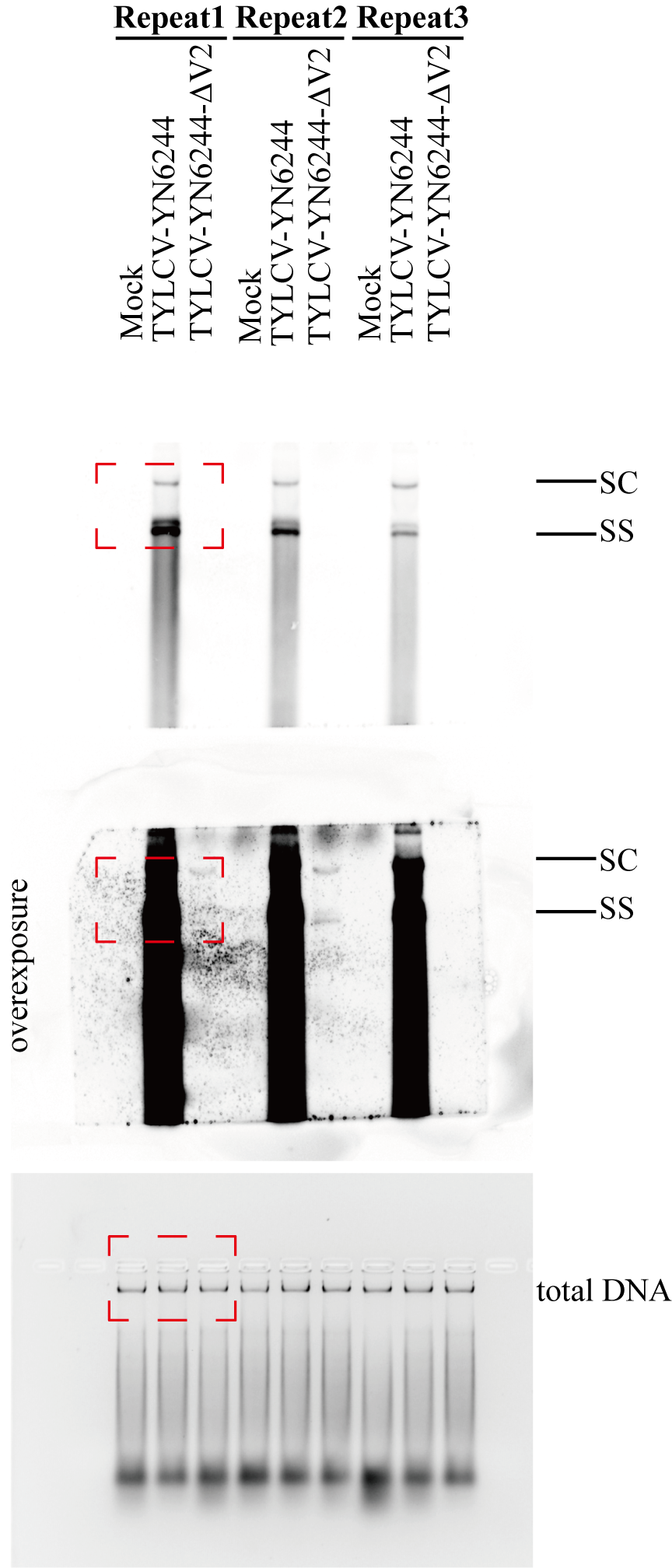
**Fig. S8** Distinct pathogenicity and impact on vsiRNA biogenesis of TYLCV-YN6244 and TYLCV-YN6244-ΔV2 in tomato, related to Fig. 4. The red dashed box corresponding to Fig. 4B. Southern blot analyses of viral genomic DNA in Micro-Tom tomato at 21 dpi after infection with TYLCV-YN6244 or TYLCV-YN6244-ΔV2. Total DNA was used as a loading control. SC, supercoiled double-stranded DNA; SS, single-stranded DNA.

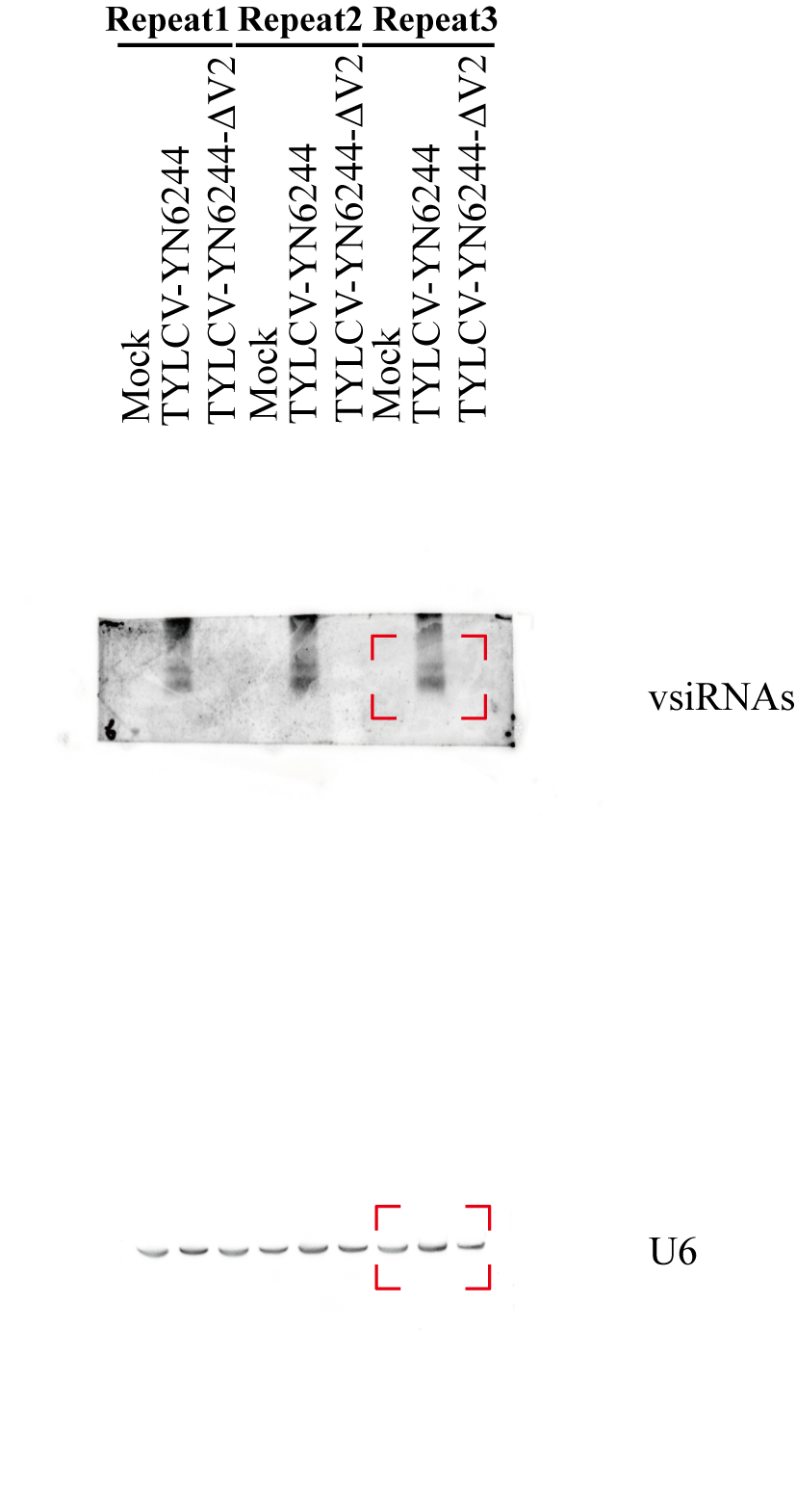
**Fig. S9** Distinct pathogenicity and impact on vsiRNA biogenesis of TYLCV-YN6244 and TYLCV-YN6244-ΔV2 in tomato, related to Fig. 4. The red dashed box corresponding to Fig. 4C. Northern blot analyses of vsiRNA accumulation in Micro-Tom tomato at 21 dpi after infection with TYLCV-YN6244 or TYLCV-YN6244-ΔV2. U6 RNA was used as a loading control.
