## Supplementary Table for "Identification and molecular characterization of a novel TYLCV isolate breaking bred-resistance to threaten tomato cultivar"

**Table S1** TYLCV-YN6244 genome sequence.

ACCGGATGGCCGCGCTTTTTCCTTTTATGTGGTCCCCACGAGTGTTCCACAGACGTCACTGTCAACCAATCAAATTGCATCCTCAAACGTTAGATAAGTGTTCATTTGTCTTTATATACTTGGTCCCCAAGTAGTTTGTCTTGCACTATGTGGGATCCACTTCTAAATGAATTTCCTGAATCTGTTCACGGATTTCGTTGTATGTTAGCTATTAAATATTTGCAGTCCGTTGAGGAAACTTACGAGCCCAATACATTGGGCCACGATTTAATTAGGGATCTTATATCTGTTGTAAGGGCCCGTGACTATGTCGAAGCGACCAGGCGATATAATCATTTCCACGCCCGTCTCGAAGGTTCGCCGAAGGCTGAACTTCGACAGCCCATACAGCAGCCGTGCTGCTGTCCCCATTGTCCAAGGCACAAACAAGCGACGATCATGGACGTACAGGCCCATGTACCGGAAGCCCAAAATATACAGAATGTATCGAAGCCCTGACGTTCCTCGTGGATGTGAAGGCCCATGTAAAGTCCAGTCTTATGAGCAACGGGATGATATTAAGCATACTGGTATTGTTCGTTGTGTTAGTGATGTTACTCGTGGATCCGGAATTACTCACAGAGTGGGTAAGAGGTTCTGTGTTAAATCGATATATTTTTTAGGTAAAGTCTGGATGGATGAAAATATTAAGAAGCAGAATCACACTAATCAGGTCATGTTCTTCTTGGTTCGTGATAGAAGGCCCTATGGAAACAGCCCAATGGATTTTGGACAGGTTTTTAATATGTTCGATAATGAGCCCAGTACCGCAACCGTGAAGAATGATTTGCGGGATAGGTTTCAAGTGATGAGGAAATTTCATGCTACAGTTATTGGTGGGCCCTCTGGAATGAAGGAACAGGCATTAGTTAAGAGATTTTTTAGAATTAACAGTCATGTAACTTATAATCATCAGGAGGCAGCCAAGTATGAGAACCATACTGAAAACGCCTTGTTATTGTATATGGCATGTACGCATGCCTCTAATCCAGTGTATGCAACTATGAAAATACGCATCTATTTCTATGATTCAATATCAAATTAATAAAATTTATATTTTATATCATGAGTTTCTGTTACATTTATTGTGTTTTCAAGTACATCATACAATACATGATCAACTGCTCTGATTACATTGTTAATTGAAATTACACCAAGACTATCTAAATACTTAAGAACTTGATATCTAAATACTCTTAAGAAACGACCAGTCTGAGGCTGTAATGTCGTCCAAATTCGGAAGTTGAGAAAACATTTGTGAATCCCCAATACCTTCCTGATGTTGTGGTTGAATCTTATCTGAATGGAAATGATGTCGTGGTTCATTAGAAATGGCCTCTGGCTGTGTTCTGTTATCTTGAAATAGAGGGGATTGTTTATCTCCCAGATAAAAACGCCATTCTCTGCTTGAGGAGCAGTGATGAGTTCCCCTGTGCGTGAATCCATGATTGTTGCAGTTGATGTGGAGGTAGTATGAGCAGCCACAGTCTAGGTCTACACGCTTACGCCTTATTGGTTTCTTCTTGGCTATCTTGTGTTGGACCTTGATTGATACTTGCGAACAGTGGCTCGTAGAGGGTGACGAAGGTTGCATTCTTGAGAGCCCAATTTTTCAAGGATATGTTTTTTTCTTCGTCTAGATATTCCCTATATGAGGAGGTAGGTCCTGGATTGCAGAGGAAGATAGTGGGAATTCCCCCTTTAATTTGAATGGGCTTCCCGTACTTTGTGTTGCTTTGCCAGTCCCTCTGGGCCCCCATGAATTCCTTGAAGTGCTTTAAATAATGCGGGTCTACGTCATCAATGACGTTGTACCAGGCATCATTACTGTACACCTTTGGGCTTAGGTCTAGATGTCCACATAAATAATTATGTGGGCCTAGAGACCTGGCCCACATTGTTTTGCCTGTTCTGCTATCACCCTCAATGACAATACTTATGGGTCTCCATGGCCGCGCAGCGGAAGATACGACGTTCTCGGCGACCCACTCTTCAAGTTCATCTGGAACTTGATTAAAAGAAGAAGAAAGAAATGGAGAAACATAAACTTCTAAAGGAGGACTAAAAATCCTATCTAAATTTGAACTTAAATTATGAAATTGTAAAATATAGTCCTTTGGGGCCTTCTCTTTTAATATATTGAGGGCCTCGGATTTACTGCCTGAATTGAGTGCCTCGGCATATGCGTCGTTGGCAGATTGCTGACCTCCTCTAGCTGATCTGCCATCGATTTGGAAAACTCCAAAATCAATGAAGTCTCCGTCTTTCTCCACGTAGGTCTTGACATCTGTTGAGCTCTTAGCTGCCTGAATGTTCGGATGGAAATGTGCTGACCTGTTTGGGGATACCAGGTCGAAGAACCGTTGGTTCTTACATTGGTATTTGCCTTCGAATTGGATAAGCACATGGAGATGTGGTTCCCCATTCTCGTGGAGTTCTCTGCAAACTTTGATGTATTTTTTATTTGTTGGGGTTTCTAGGTTTTTTAATTGGGAAAGTGCTTCCTCTTTAGAGAGAGAACAATTGGGATATGTTAGGAAATAATTTTTGGCATATATTTTAAATAAACGAGGCATGTTGAAATGAATTGGTGTCCCTCAAAGCTCTATGGCAATCGGTGTATCGGTGTCTTACTTATACCTGGACACCTAATGGCTATTTGGTAATTTTGTAAAAGTACATTGCAATTCAAAATTCAAAATTCAAAAATCAAATCATTAAAGCGGCCATCCGTATAATATT

**Table S2** Coding genes and Intergenic region (IR) of TYLCV-YN6244.

| TYLCV-YN6244 | 2781nt |
| --- | --- |
| *V1* | From 308 ^th^ to 1084 ^th^ (777 nt) |
| *V2* | From 148 ^th^ to 498 ^th^ (351 nt) |
| *V3* | From 2350 ^th^ to 2583 ^th^ (234 nt) |
| *C1* | From 1542 ^th^ to 2615 ^th^ (1074 nt) |
| *C2* | From 1226 ^th^ to1633 ^th^ (408 nt) |
| *C3* | From 1081 ^th^ to 1485 ^th^ (405 nt) |
| *C4* | From 2171 ^th^ to 2464 ^th^ (294 nt) |
| *C5* | From 659 ^th^ to 862 ^th^ (204 nt) |
| IR | From 2616 ^th^ to 147 ^th^ (313 nt) |

**Table S3** Known TYLCV isolates used for phylogenetic tree analysis.

| No. | Isolate | GenBank acc. No. |
| --- | --- | --- |
| 1 | TYLCV-Alm | AJ489258 |
| 2 | TYLCV_AU_1923-2006 | KX347097 |
| 3 | TYLCV-BJ | MN432609 |
| 4 | TYLCV-SC780 | MK559473 |
| 5 | TYLCV-SDSG | HQ702862 |
| 6 | TYLCV-SH2 | AM282874 |
| 7 | TYLCV-SJZ1 | JF727878 |
| 8 | TYLCV-SX8 | JN412854 |
| 9 | TYLCV-USA | EF539831 |
| 10 | TYLCV-YN6244 | in this study |
| 11 | TYLCV-ZJ8 | AM698119 |

**Table S4** PCR analyses of systemic infection rate of TYLCV-YN6244 and TYLCV-YN6244-ΔV2 in tobacco plants. PCR analysis was conducted via primer pairs of TYLCV-YN6244-IR-F/R.

| **Infectious clone** | **Number of plants infected / total number of inoculated plants** | **Infection rate** |
| --- | --- | --- |
| TYLCV-YN6244 | 36/36 | 100% |
| TYLCV-YN6244-ΔV2 | 36/36 | 100% |

**Table S5** Primers used in this study.

| **Used for** | **Primer name** | **Sequence 5' to 3'** |
| --- | --- | --- |
| Construction of TYLCV-YN6244-V1 expression vector | pCAMBIA3301-flag-V1-F | GACGATGACAAGACGCGTCCCGGGTCGAAGCGACCAGGCGATATAATC |
|  | pCAMBIA3301-flag-V1-R | CATTATTATGGAGAAAGCTTGGATCCTTAATTTGATATTGAATCATAG |
| Construction of TYLCV-YN6244-V2 expression vector | pCAMBIA3301-flag-V2-F | GACGATGACAAGACGCGTCCCGGGTGGGATCCACTTCTAAATGAAT |
|  | pCAMBIA3301-flag-V2-R | CATTATTATGGAGAAAGCTTGGATCCTCAGGGCTTCGATACATTCTG |
| Construction of TYLCV-YN6244-V3 expression vector | pCAMBIA3301-flag-V3-F | GACGATGACAAGACGCGTCCCGGGTTCGGATGGAAATGTGCTGACC |
|  | pCAMBIA3301-flag-V3-R | CATTATTATGGAGAAAGCTTGGATCCTTATTTCCTAACATATCCCAAT |
| Construction of TYLCV-YN6244-C1 expression vector | pCAMBIA3301-flag-C1-F | GACGATGACAAGACGCGTCCCGGGCCTCGTTTATTTAAAATATATG |
|  | pCAMBIA3301-flag-C1-R | CATTATTATGGAGAAAGCTTGGATCCTTACGCCTTATTGGTTTCTTC |
| Construction of TYLCV-YN6244-C2 expression vector | pCAMBIA3301-flag-C2-F | GACGATGACAAGACGCGTCCCGGGCAACCTTCGTCACCCTCTACGAG |
|  | pCAMBIA3301-flag-C2-R | CATTATTATGGAGAAAGCTTGGATCCCTAAATACTCTTAAGAAACGAC |
| Construction of TYLCV-YN6244-C3 expression vector | pCAMBIA3301-flag-C3-F | GACGATGACAAGACGCGTCCCGGGGATTCACGCACAGGGGAACTC |
|  | pCAMBIA3301-flag-C3-R | CATTATTATGGAGAAAGCTTGGATCCTTAATAAAATTTATATTTTATA |
| Construction of TYLCV-YN6244-C4 expression vector | pCAMBIA3301-flag-C4-F | GACGATGACAAGACGCGTCCCGGGGGGAACCACATCTCCATGTG |
|  | pCAMBIA3301-flag-C4-R | CATTATTATGGAGAAAGCTTGGATCCTTAATATATTGAGGGCCTCG |
| Construction of TYLCV-YN6244-C5 expression vector | pCAMBIA3301-flag-C5-F | GACGATGACAAGACGCGTCCCGGGAAATTTCCTCATCACTTGAAAC |
|  | pCAMBIA3301-flag-C5-R | CATTATTATGGAGAAAGCTTGGATCCTTAGGTAAAGTCTGGATGGATG |
| Construction of TYLCV-YN6244-C5-1 expression vector | pCAMBIA3301-flag-C5-1-F | GACGATGACAAGACGCGTCCCGGGGGCCTGTACGTCCATGATCGTC |
|  | pCAMBIA3301-flag-C5-1-R | CATTATTATGGAGAAAGCTTGGATCCTTAATTAGGGATCTTATATCTG |
| Construction of pBinplus-0.5-unit TYLCV-YN6244 vector | TYLCV-YN6244-0.5 unit-F | ATGCTACAGTTATTGGTGGGC |
|  | TYLCV-YN6244 -0.5 unit-R | CCCAAGCTTGCCCACCAATAACTGTAGCAT |
| Construction of pBinplus-1-unit TYLCV-YN6244 vector | TYLCV-YN6244 -1 unit-F | CGAGCTCTTAGCTGCCTGAATGTTCGG |
|  | TYLCV-YN6244 -1 unit-R | CGAGCTCAACAGATGTCAAGACCTACGTG |
| Construction of pBinplus-0.5-unit TYLCV-YN6244-ΔV2 vector | TYLCV-YN6244-ΔV2-0.5 unit-F | CTTACGAGCCCAATACATTG |
|  | TYLCV-YN6244-ΔV2-0.5 unit-R | GTGCAAGACAAACTACTTGG |
| Construction of pBinplus-1-unit TYLCV-YN6244-ΔV2 vector | TYLCV-YN6244-ΔV2-1 unit-F | ATGTTAGCTATTAAATATTTGC |
|  | TYLCV-YN6244-ΔV2-1 unit-R | GTGCAAGACAAACTACTTGG |
| Detection TYLCV-YN6244 and TYLCV-YN6244-ΔV2 | TYLCV-YN6244-IR-F | GTTGAAATGAATTGGTGTCC |
|  | TYLCV-YN6244-IR-R | ATGGCAAGACAAACTACTTG |
| qPCR analysis of viral DNA accumulation | qTYLCV-C1-F | TACTTTGTGTTGCTTTGCCAG |
|  | qTYLCV-C1-R | CCAGGTCTCTAGGCCCACATA |
| Probe labeling with PCR for Southern blot and Northern blot | Probe-F | GACGATGACAAGACGCGTCCCGGGTCGAAGCGACCAGGCGATATAATC |
|  | Probe-R | CATTATTATGGAGAAAGCTTGGATCCTTAATTTGATATTGAATCATAG |
| Identification *Ty*-1 in Zuanhong No. 5 | *Ty-1*-F | CAACAGCAATGTACCTGGTCAG |
|  | *Ty-1*-R | CTGTGGCATACGTTGGTGACAC |
| Identification *Ty*-2 in Baxi | *Ty-2*-F | TGGCTCATCCTGAAGCTGATAGCGC |
|  | *Ty-2-*R | AGTGTACATCCTTGCCATTGACT |
